# Chronic modulation of human memory and thalamic-hippocampal theta activities

**DOI:** 10.1101/2022.12.21.521275

**Authors:** Victoria S. Marks, Michal Lech, Nicholas M. Gregg, Vladimir Sladky, Filip Mivalt, Dan P. Crepeau, Jaromir Dolezal, Eva Alden, Brian N. Lundstrom, Bryan Klassen, Steven A. Messina, Benjamin H. Brinkmann, Kai J. Miller, Jamie J. Van Gompel, Vaclav Kremen, Gregory A. Worrell, Michal T. Kucewicz

**Affiliations:** Mayo Clinic Graduate School of Biomedical Sciences, Mayo Clinic, Rochester, Minnesota, USA; Bioelectronics, Neurophysiology, and Engineering Laboratory, Department of Neurology, Mayo Clinic, Rochester, Minnesota, USA; BioTechMed Center, Multimedia Systems Department, Faculty of Electronics, Telecommunications and Informatics, Gdansk University of Technology, Gdansk, Poland; Faculty of Biomedical Engineering, Czech Technical University in Prague, Prague, Kladno, Czech Republic; Department of Biomedical Engineering, Faculty of Electrical Engineering and Communication, Brno University of Technology, Brno, Czech Republic; Czech Institute of Informatics, Robotics, and Cybernetics, Czech Technical University in Prague, Prague, Czech Republic; Division of Neurocognitive Disorders, Department of Psychiatry and Psychology, Mayo Clinic, Rochester Minnesota, USA; Department of Radiology, Mayo Clinic, Rochester, Minnesota, USA; Department of Neurosurgery, Mayo Clinic, Rochester, Minnesota, USA

## Abstract

Electrical stimulation is a powerful therapeutic tool for treating neurologic and neuropsychiatric disorders. Sensing and modulating electrophysiological biomarkers of memory over extended timescales is necessary for tracking and improving memory in humans. Here, we describe results from humans in their natural home environments using a novel, investigational system enabling chronic stimulation and multi-channel recording of anterior thalamic and hippocampal local field potentials during memory tasks. Four people with focal epilepsy performed a free recall verbal memory task repeatedly for up to fifty months on a touch-screen device with wireless signal acquisition with electrophysiology and behavioral data streaming to a cloud environment. Anterior thalamic-hippocampal spectral activities in the theta frequency range were found to correlate with memory processing, to predict task performance, and to be modulated by deep brain stimulation. Our results provide a new biomarker-based technology for chronic remote tracking of memory performance and modulation of the associated neural activities.

**One Sentence Summary:** Electrical stimulation in the anterior thalamic nuclei modulates theta frequency activities and improves human verbal memory performance chronically.

## Introduction

Since the clinical success of Deep Brain Stimulation (DBS) therapy for treating movement disorders, it has been applied in a range of neurological and neuropsychiatric conditions*(1)*, including obsessive compulsive disorder*(2)*, Tourrette’s syndrome*(3)*, and depression*(4, 5)*. Cognitive functions like declarative memory have been modulated by DBS and less invasive stimulation methods with inconsistent results*(6–10)* not replicated across studies*(11, 12)* and which focused mainly on acute task performance effects. Clinical trials for treating memory deficits in neurodegenerative disorders, for example, hippocampal fornix DBS in Alzheimer’s disease, have so far had limited success *(13–16)*. Another trial of DBS in the nucleus basalis of Meynert for Lewi body dementia *(17, 18)* also obtained limited individual effects*(19)*, despite promising results from preceding studies *(20, 21)*. Thus there is a lack of conclusive evidence for robust positive effects on declarative memory performance.

A theory-driven approach based on electrophysiological biomarkers of neural activities is one plausible strategy for DBS studies of cognitive functions. Instead of modulating behavioral outcomes indirectly via an unknown underlying neurophysiology, it may be more feasible to modulate neural activities based on biomarker feedback to affect the desired outcomes*(22)*. This approach requires continuous recordings and analysis of the biomarker activities supporting memory processing. In our previous studies, we developed technology for concurrent telemetric recording and stimulation from an implantable device*(23)* with signal acquisition and processing distributed to external devices and cloud environments *(24–26)*. This provided continuous tracking of neural activities (in this case, seizures) and behavioral states in implanted patients throughout their daily lives *(26, 27)* to inform DBS parameter adjustment according to the analyzed biomarker activities. Such technologies were successfully implemented in the management of epilepsy *(28–30)* and Parkinson’s disease *(31, 32)*, where the neural biomarkers of pathophysiological activities are well studied and validated.

In the case of memory and cognitive functions, the choice of specific neurophysiological biomarkers to target are less clear. In Parkinson’s disease, beta neural oscillations that are associated with motor deficits have been successfully used as a biomarker for adaptive DBS modulation *(33–35)*. In this example, movement dysfunction is treated by adaptively adjusting the parameters and timing of DBS in response to pathological bursts of beta activity in the affected brain regions. With regard to cognition, we can track and automatically identify general states of sleep and wakefulness*(36)*, but more specific states supporting successful memory formation are difficult to classify *(37, 38)*, with multiple possible electrographic biomarkers *(39, 40)*. Prior studies suggest that biomarker-driven stimulation results in a similar magnitude of an effect on memory performance as open-loop stimulation without feedback from recorded neural activity *(41, 42)*. Chronic tracking and validation of the biomarker activities for memory and cognitive functions across longer time scales is needed for new applications of adaptive DBS.

Here, we used a DBS system for continuous chronic recording and stimulation bilaterally in the anterior nucleus of the thalamus and in the hippocampus. The former is a target for treatment of seizures *(43, 44)* and, more recently, has been proposed as a target for cognitive functions *(45–50)*. The latter has been associated with mixed effects of acute DBS on memory performance *(11, 12, 51)*. In this study, we developed a new system for chronic tracking of memory performance and modulation based on electrophysiological biomarkers of neural activities. Cognitive improvements after long-term thalamic stimulation have been observed *(52, 53)*, and we have recently demonstrated a long-term positive advantage of low-frequency over high-frequency anterior thalamic DBS on verbal memory performance*(54)*. Since none of the previous studies continuously recorded anterior thalamic activity during cognitive tasks, this is the first test of the role of anterior thalamic-hippocampal neural activity in modulation of memory processing over months and years of regular task performance remotely from home. We hypothesized that DBS of this circuit modulates the anterior thalamic-hippocampal activity and the associated verbal memory performance chronically. Our goal was to evaluate potential chronic therapeutic modulation of memory functions resulting from a brain stimulation approach for epilepsy management in real-life environments.

## Results

In the four patients implanted with the Medtronic Summit RC+S™ system, we collected a total of 270 weeks (mean 67.5 weeks per patient) of continuous ANT and Hc local field potential (LFP) recordings during daily life activities, periodically testing performance on a free recall verbal memory task (Table 1). The task sessions were performed by the patients in their homes with remote telephone assistance to guide and provide encouragement. These were performed regularly up to once a week for a maximum period of two years. Overall, behavioral and LFP data were collected from 117 task sessions (min. 9, max. 51) corresponding to a total of 1755 word lists (each list comprising 12 words). Three of the patients showed >70% of all LFP data successfully streamed*(54)*, and the streaming rate from the task sessions was >95% for all four patients. During the task performance, LFP data was streamed from the same set of electrode leads localized bilaterally to the anterior nuclei of the thalamus and hippocampi (Fig. 1a) with stimulation paused. All patients were able to initiate and complete the task sessions independently on the touch-screen devices with remote supervision (Fig. 1b). The LFP signals were synchronized with the task events with <50ms delay and without significant data loss. We confirmed that task-induced changes in the spectral power and coherence were time locked to the task events (Fig. 1c-d), verifying accurate synchronization, and were predominantly limited to the theta frequency range (4-8 Hz). Hence, analysis focused on this frequency band, with other bands summarized in the supplementary material.

**Table 1.** Since implantation, patients have received bilateral continuous ANT stimulation interrupted by regular performance of the verbal memory task with ANT and Hc sensing. Gold standard seizure counts, number of the task sessions, trials and hours are summarized in the table. Note that each task session comprises 15 lists of 12 words. Note: M2 withdrew from the study before implantation. ANT - anterior nucleus of the thalamus; DBS - deep brain stimulation.

| Patient ID | Months since<br>implantation | Seizures/month | Task<br>sessions | Word<br>lists | Task<br>sessions<br>with<br>robust<br>data | Word lists<br>within tasks<br>with robust<br>data |
| --- | --- | --- | --- | --- | --- | --- |
| M1 | 28 | 20.6 | 51 | 765 | 38 | 570 |
| M3 | 13 | 293.3 | 21 | 315 | 18 | 270 |
| M4 | 9 | 26.7 | 9 | 135 | 9 | 135 |
| M5 | 13 | 4.4 | 36 | 540 | 31 | 465 |

**Fig. 1.**
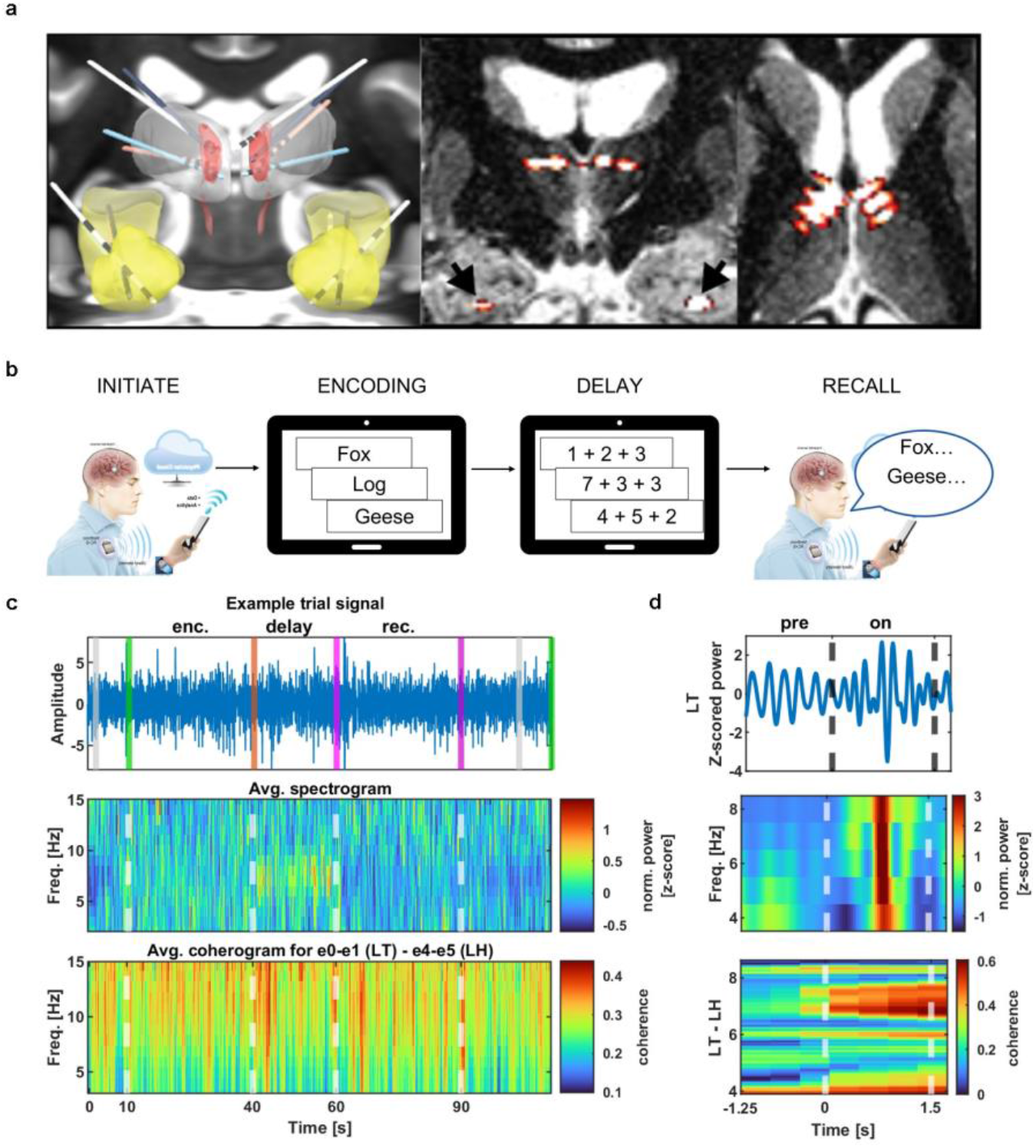
Continuous chronic recordings of anterior thalamic and hippocampal LFP are recorded during remote memory task performance. (**a**) Electrode locations from all patients are shown superimposed on coregistered CT and MRI, with hippocampi highlighted in yellow and anterior nuclei of the thalamus highlighted with red color in the left panel. Middle and right panels show an example localization in patient M1 implantation after coregistration with pre-implant MRI. Notice that all electrode contacts are within the ANT and Hc bilaterally. (**b**) Tasks were remotely initiated by the investigator while on the telephone with the patient. The encoding phase comprised 12 successive word presentations on the screen for each list trial. The delay phase lasted 20 s when the patient solved simple algebraic equations. During the recall phase, the patient stated as many words from the list as they could remember. (**c**) Example signal from one trial of patient M5 with colored lines marking task events: gray – trial start, green – encoding start, orange – delay start, purple – recall phase onset and offset. An average spectrogram and coherogram from all 31 trials of patient M5 are presented. Notice how the theta activity increases during the delay phase and how coherence increases just after the beginning of the delay phase. (**d**) Example theta-filtered signal from the left thalamus during a recalled word by patient M5 is presented as a spectrogram of ANT activity and coherogram from a pair of left thalamus–left hippocampus (LT-LH) leads. “0” onset marks the moment of the word display, preceded by 1-sec blank screen period. The word disappears from the screen after 1.5 s. Notice the increase in the left thalamic power and the thalamic-hippocampal coherence induced by the word display event.

### Anterior thalamic-hippocampal theta activities are associated with verbal memory

We first investigated whether the LFP activity in ANT and Hc was modulated by the task performance. Differences in normalized spectral power in and coherence between these brain areas were compared between the encoding, delay, and recall phases of the task (Fig. 2a). In the left hemisphere, power of theta frequency activity (4-8 Hz) was significantly higher during the delay phase than the recall phase in the hippocampus (Kruskal-Wallis test, *χ*^2^=9.27, p=0.009; Tukey-Kramer post-hoc test, p=0.007) and anterior thalamus (Kruskal-Wallis test, *χ*^2^=6.04, p=0.049; Tukey-Kramer post-hoc test, p=0.038). In the right hemisphere, theta power was significantly higher during the encoding phase than the recall phase in the hippocampus (Kruskal-Wallis test, *χ*^2^=8.77, p=0.013; Tukey-Kramer post-hoc test, p=0.009) and thalamus (Kruskal-Wallis test, *χ*^2^=7.04, p=0.03; Tukey-Kramer post-hoc test, p=0.02). The increased theta power (but not coherence; see Supplementary Fig. S1-3 for other frequency bands) during the encoding and delay phases, when the cognitive load was the highest, suggests a role of ANT-Hc theta activities in formation and maintenance of memory items.

**Fig. 2.**
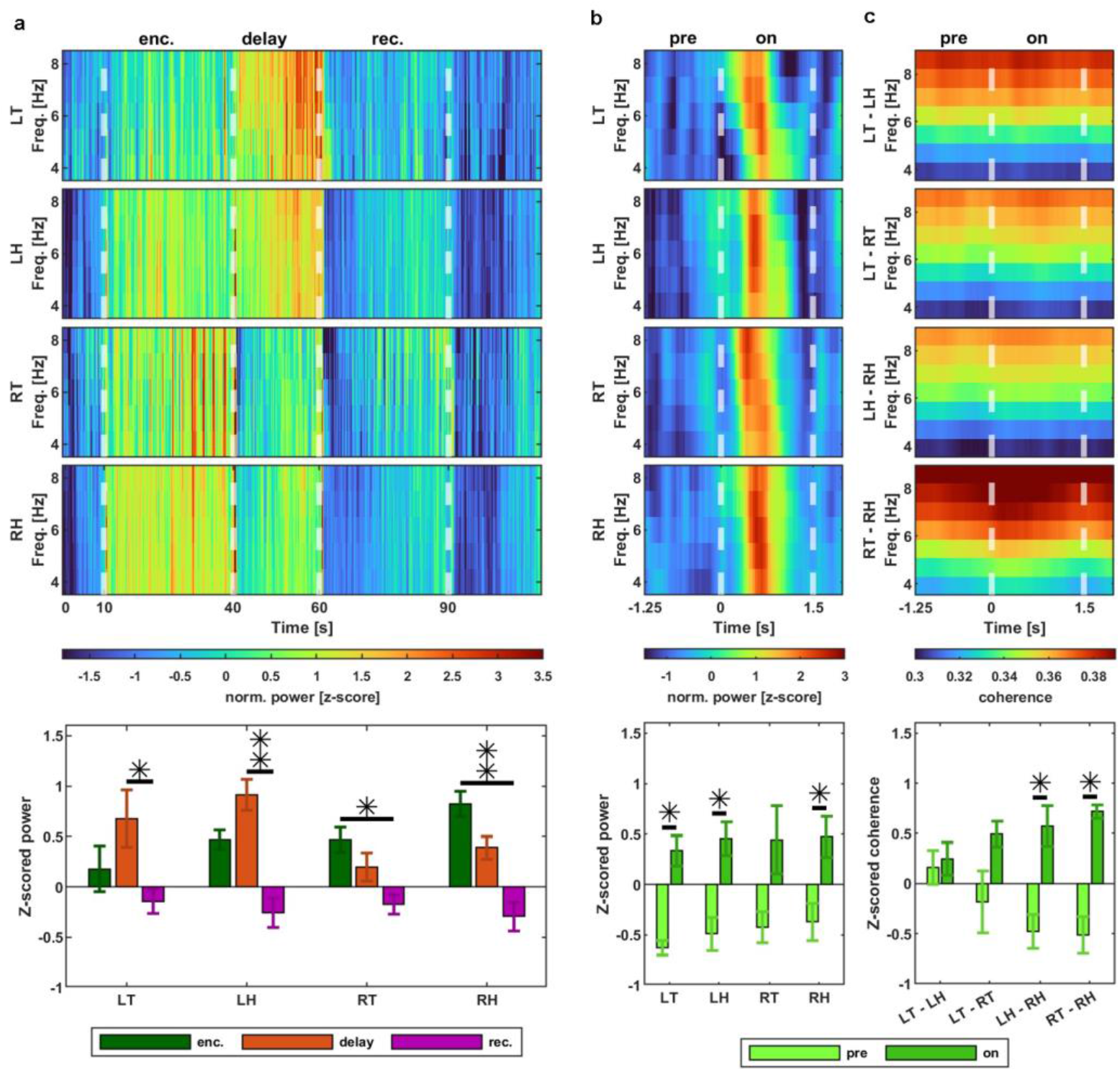
Anterior thalamic-hippocampal neural activities in the theta frequency band are modulated by verbal memory task. (**a**) The grand-average theta power (all patients) in the left thalamus and the left hippocampus was higher during the delay than during the recall (rec.) phase; in the right thalamus and hippocampus it was higher during the encoding (enc.) compared to the recall phase. (**b**) The theta power was increased during word presentation for encoding (1s after onset) compared to immediately before (1s preceding onset) in the left thalamus and both hippocampi. (**c**) The grand-average amplitude of the theta coherence was significantly greater during the word presentation than before the word onset between LH-RH and RT-RH lead pairs.

Next, we looked at modulation of theta power and coherence on the discrete level of individual word (encoding) and equation (delay) presentation events. We found that theta power was significantly higher, i.e. induced, during word presentation relative to the baseline before presentation (Fig. 2b) in the left thalamus (LT) and in both left (LH) and right (RH) hippocampi (Kruskal-Wallis test; *χ*^2^=5.33, p=0.02; *χ*^2^=5.33, p=0.02; *χ*^2^=4.08, p=0.04; respectively) but not in right thalamus (RT). Likewise, alpha power was significantly induced during word presentation in LT and LH (Kruskal-Wallis test; *χ*^2^=5.33, p=0.02; *χ*^2^=4.08, p=0.04; respectively; Supplementary Fig. S1b).

We also found an analogous induction of theta coherence (Fig. 2c) between the LH-RH and RT-RH lead pairs (Kruskal-Wallis test, *χ*^2^=5.33, p=0.02, in both cases). Conversely, coherence in the gamma band decreased at word presentation between LT-LH (Kruskal-Wallis test; *χ*^2^=4.08, p=0.04; Supplementary Fig. S3c). There were no significant changes in theta power or coherence in response to equation presentation (Supplementary Fig S4a) - these were specific to memory encoding phase. The equation display events induced beta and gamma power in LT and RT (Kruskal Wallis test, *χ*^2^=5.33, p=0.02, in each case) and induced gamma coherence between LH-RH and RT-RH (Supplementary Fig. S4c-d; Kruskal Wallis test, *χ*^2^=4.08, p=0.04 *χ*^2^=5.33, p=0.02). We conclude that the theta activities were selectively induced by the memory encoding events but not the distractor events.

### Theta activities during word encoding predict subsequent recall

We hypothesized that the theta power and coherence changes could be used to predict successful recall, i.e. show subsequent memory effect*(55)*. First of all, we found that task-induced changes in the theta power were different between trials with subsequently recalled and forgotten words (Fig. 3a). Theta power was significantly induced by the recalled (but not the forgotten) word presentation in RT and RH (Kruskal-Wallis, *χ*^2^=4.08, p=0.04; *χ*^2^=5.33, p=0.02, respectively). In contrast, theta power was significantly induced by the forgotten (but not the recalled) word presentation in LT and LH (Kruskal-Wallis, *χ*^2^=5.33, p=0.02 for both). Similar trial-specific power inductions were found in the other bands: alpha and gamma power were induced by the forgotten words in the LH (Kruskal-Wallis, *χ*^2^=5.33, p=0.02; *χ*^2^=4.08, p=0.04, respectively; Supplementary Fig. S5a and S7a), and alpha power was induced by the recalled words in the RH (Kruskal-Wallis, *χ*^2^=4.08, p=0.04; Supplementary Fig. S5a). Importantly, subsequent memory effect between the recalled and forgotten trials was only found in LT and LH theta power, which was significantly lower before the onset of the forgotten words (Kruskal-Wallis, LT: *χ*^2^=5.33, p=0.02; LH: *χ*^2^=4.08, p=0.04). In other words, suppressed theta power in left ANT-Hc before word onset predicted unsuccessful memory encoding.

**Fig. 3.**
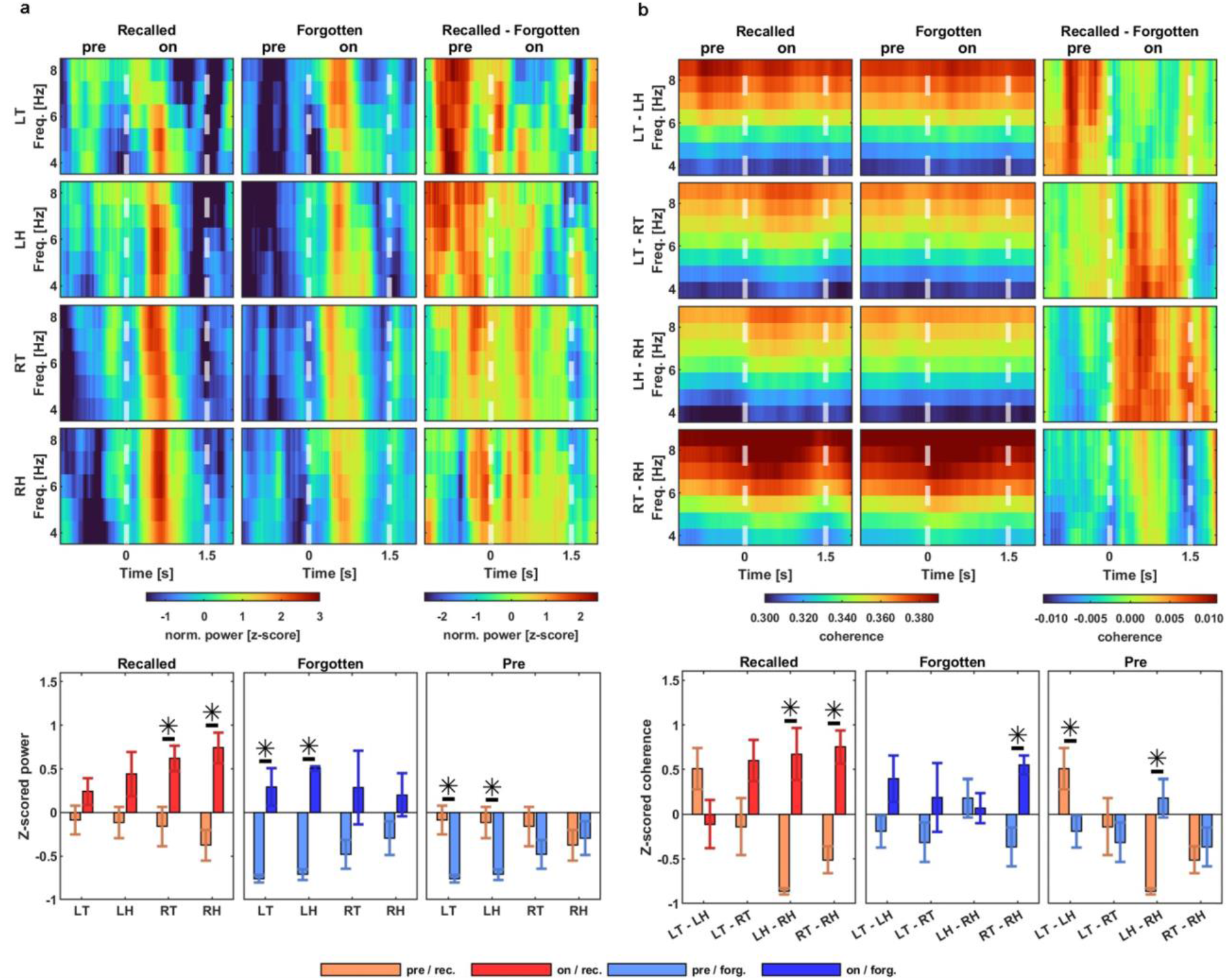
Theta power and coherence across the anterior thalamic nuclei and the hippocampi are associated with successful memory processing. (**a**) Theta power was significantly induced by the display of subsequently recalled words in the right thalamus and hippocampus (left panels). Subsequently forgotten words, on the other hand, induced a significant theta power response in the left thalamus and hippocampus (middle panels). Subsequent memory effect was observed in the left thalamus and hippocampus where theta power was significantly lower before the presentation of the forgotten compared to the recalled words. (**b**) Analogous analysis for coherence revealed significantly induced connectivity between both hippocampi on the recalled word trials (left panels) and between the right thalamus and hippocampus on both recalled and forgotten word trials (left and middle panels). The subsequent memory effect was again observed between the left thalamus and hippocampus (LT-LH) with coherence significantly greater before presentation of the recalled words, and between both hippocampi (LH-RH) where the theta coherence was significantly lower before presentation of the recalled words. Notice that both thalamic and hippocampal power and coherence predict subsequent recall in the period immediately preceding word encoding.

The importance of the left ANT-Hc theta interactions for verbal memory encoding was also reflected in theta coherence (Fig. 3b). Trial-specific changes were observed between the hippocampi (LH-RH) during the recalled words (Kruskal-Wallis, *χ*^2^=5.33, p=0.02) and between RT-RH in both recalled and forgotten trial types (Kruskal-Wallis, *χ*^2^=5.33, p=0.02, in both cases). The subsequent memory effect, however, was specific to LT-LH, showing greater theta coherence immediately before presentation of the recalled compared to the forgotten words (Kruskal-Wallis, *χ*^2^=4.08, p=0.04). A reverse memory effect was found between both hippocampi (LH-RH), i.e., theta coherence was significantly lower before presentation of recalled words than forgotten (Kruskal-Wallis, *χ*^2^=5.33, p=0.02). All in all, only an increase in LT-LH theta coherence and a decrease in the LH-RH theta coherence predicted successful memory recall.

We conclude that theta power and coherence show consistent differences between successful and failed memory encoding, especially in the period immediately preceding stimulus presentation. The pattern of task-induced theta power (Fig. 3a) and coherence (Fig. 3b) was overlapping.

### Chronic memory performance is reflected by the thalamic-hippocampal theta activities

Having established that the theta activities within and between ANT and Hc predict successful recall of verbal memory acutely, we investigated whether they would predict general memory performance chronically across multiple sessions. We previously found that bilateral ANT stimulation with low frequency current had a positive effect on the task performance compared to high frequency stimulation*(54)*. We first confirmed the positive effect in two patients with the longest track of recordings. We found 43-46% improvement in the mean rate of words recalled between sessions performed during the high frequency (145 Hz) and the low frequency (2 Hz) stimulation modes (Fig. 4a; M1: from 0.358 to 0.450, M5: from 0.251 to 0.308).

**Fig. 4.**
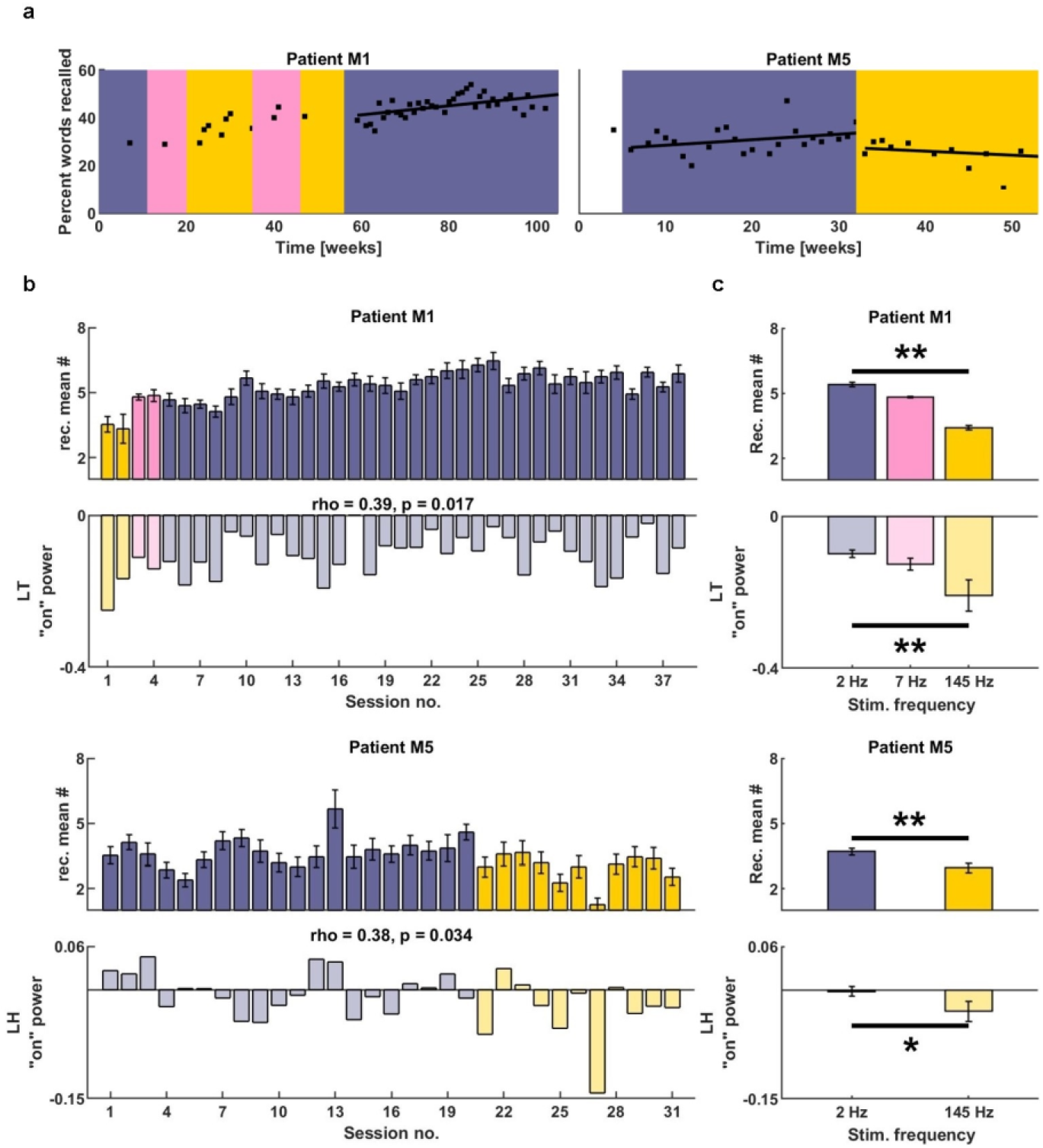
Longitudinal memory performance and the effect of DBS are reflected in the thalamic-hippocampal theta activities. (**a**) Individual patient examples show progressive improvement in memory performance with low-frequency (2 Hz) ANT stimulation mode (violet background) and gradual deterioration in performance with high-frequency (145 Hz) DBS (yellow). Pink indicates 7 Hz DBS, and white background indicates no chronic DBS. (**b**) Mean memory performance is correlated with the mean theta power at word onset (background DBS color-coded as in (**a**)). (**c**) Patients recalled fewer words and showed less theta power at word onset in the left thalamus (M1) and the left hippocampus (M5) during the high-frequency stimulation mode of chronic DBS. Note: a subset of sessions from (**a**) with robust data was used in (**b**) and (**c**).

This behavioral improvement in task performance was correlated across time with the theta neural activities induced during word encoding (Fig. 4b) - similar to the acute trial-level memory effects but on the global level of sessions. There was a significant correlation of mean session performance with the LT “pre-onset” theta power in patient M1 (Spearman, r=0.39, p=0.017; Fig. 4b), with the LH “pre” and “onset” theta power in patient M5 (Spearman, r=0.45, p=0.011, and r=0.38, p=0.034, respectively; Fig. 4b and Supplementary Fig. S8), and with “pre-onset” RT theta power in patient M5 (Spearman, r=0.43, p=0.016; Supplementary Fig. S8). The other two patients had insufficient data for this longitudinal correlation with the two stimulation modes. In all these cases, higher power of the theta activities during memory encoding indicated good performance in the memory task.

Finally, the modulated theta activities reflected the effect of ANT stimulation on memory performance. Patient performance gradually improved with the low-frequency mode of stimulation (Pearson; M1: r=0.56, p=0.0002; M3: r=0.63, p=0.0039). In general, patients recalled more words when in the low-frequency stimulation mode (permutation test; p=0.002 for patient M1, p=0.004 for patient M5; Fig. 4c). Task performance during this stimulation mode revealed significantly higher theta power in LT of patient M1 (permutation test, p=0.008), and in LH of patient M5 (permutation test, p=0.04; Fig. 4c) compared to the high-frequency stimulation mode. This chronic modulatory effect of ANT stimulation carried over from outside of memory performance, since it was halted during the tasks.

## Discussion

Our study provides the first demonstration of continuous chronic deep-brain recordings from the anterior nucleus of the thalamus and the hippocampus during regular memory tasks performed remotely from real-life, home environments. This is a major technical advancement from the recent demonstration of deep-brain hippocampal recordings with ambulatory patients performing a spatial navigation task in a real-life laboratory environment*(56)*. Taking the human intracranial recordings from a laboratory setup with limited time to collect data to the outside environment with continuous unlimited data streaming and online cloud analysis is a new milestone for next-generation brain stimulation technologies. The technology we employed in this study was developed from its original applications for adaptive therapeutic stimulation for epilepsy *(24, 25)* and Parkinson’s disease*(31)*. Thanks to the long-term recording and analysis distributed to remote devices and cloud computations, it opens a wide range of opportunities for studying and treating memory and cognitive functions with tracking of behavioral states of sleep and wakefulness*(27)*, circadian and ultradian cycles *(29, 57)*, as well as mood and cognitive assessments*(58)*.

Here, we prototyped a behavioral paradigm for regular repeated longitudinal testing of short-term verbal memory that is resistant to training and habituation effects to show: 1) correlation of ANT-Hc neural activities with acute and chronic memory performance, 2) co-modulation of the theta activities and behavioral accuracy by ANT stimulation, and 3) large and lasting stimulation-induced improvement in verbal memory and specific theta activities. Altogether, we provide a robust technological tool and a proof-of-concept for long-term assessment of verbal memory performance and chronic modulation of the underlying neural activities. The study was limited to one patient population with particular brain pathophysiology and just one memory function. Future development of accelerometry, gaze-tracking and precise positioning will extend applications to other diseases and functions like navigation and spatial memory.

Our results show that the anterior thalamic-hippocampal activities of the Papez circuit are involved in verbal memory processing. Previously, this circuit has been implicated in spatial episodic memory, but accumulating evidence suggests non-spatial roles for this circuit, including emotional affect and memory guided attention *(45, 46, 48, 49, 59)*. Anterior nucleus of the thalamus was proposed as a core component of the episodic memory system*(47)*. Here, we demonstrated that theta activity in this circuit is involved with verbal memory processing. Firstly, theta power and coherence were induced by presentation of words for encoding. Secondly, these induced theta activities were different during successful memory trials with subsequently recalled words, showing memory effects selectively in the left anterior thalamus and between both hippocampi. The memory effects were anatomically lateralized, pointing to the typically language-dominant left hemisphere. Finally, the same theta activities induced at the onset of words for memory encoding reflected accuracy of task performance longitudinally across multiple sessions. In other words, greater theta activity prior to word onset was associated with better verbal memory performance. Even though the activities were not the same and limited to a single anatomical site in every patient, they provide electrophysiological biomarkers of memory performance within a consistent range of frequencies in a distinct anatomical circuit. The importance of theta activities likely extends beyond this specific function to other non-verbal domains, including navigation and spatial memory*(60)*. Theta activities at distinct frequencies and at particular sites of the Papez circuit may play unique roles in spatial and non-spatial memory functions, e.g. with greater involvement of the right or the left hemisphere, respectively.

We found most of the memory-related theta activities occurred immediately before word presentation for encoding. Magnitude of both theta power and theta coherence between LT-LH was predictive of successful memory formation during attentive anticipation for the words to be presented on the screen. This suggests an underlying process related to attention and preparation for stimulus encoding - critical for memorization and subsequent recall. This preparatory activity is observed at all levels of neural activities starting from single neuron firing, as recently shown in the human hippocampus and termed “attention to encoding*(61)*,” all the way to EEG theta oscillations in the thalamic nuclei *(62, 63)*. The latter showed that theta power recorded immediately before image presentation in the right anterior and mediodorsal thalamic nuclei predicted subsequent recall. In our study, the memory effect was lateralized to the left thalamus given that verbal stimuli were used instead of images. We found theta activity was also enhanced during the delay phase of the task when the remembered items must be sustained against distractor math equations, which also points to an attention-related function (i.e. working memory). Both these neural activities and the pattern of anatomical connectivity to the anterior prefrontal cortical areas are congruent with a higher-order executive function of the anterior thalamic nuclei related to memory and attention *(45–50, 64)*. A hotspot of the theta activities in the left anterior prefrontal cortex in the same task was recently identified*(40)*, confirming the highest magnitude of the memory effect immediately before word presentation in the cortical area directly connected to the anterior thalamic nuclei.

A strong predictive memory signal that starts even before the actual stimulus encoding presents an ideal biomarker for therapeutic interventions targeting cognition using DBS. Such biomarkers not only predict the possible outcomes of subsequent memory encoding but can provide feedback triggers for precise timing of brain stimulation in a closed-loop of LFP sensing and modulation. Previous attempts at direct electrical brain stimulation to enhance stimulus encoding identified a successful target in the lateral temporal cortex, resulting in approx. 15% improvement in performance *(41, 42)*. In this study, we observed up to 46% improvement in the same verbal memory task with continuous adaptive DBS in the anterior nucleus of the thalamus. These results offer a potentially more effective strategy and/or new anatomical targets for therapeutic interventions for memory and cognitive deficits. In contrast to the widespread dispersed cortical networks, the thalamic nuclei connected to these networks are compact and condensed in confined volumes of neural tissue, making them suitable for local interventions like DBS to effectively modulate larger thalamocortical networks*(65)*. Future studies will determine the exact role of the ANT theta activities in memory and cognitive functions, employing closed-loop and long-term adaptive stimulation paradigms.

The goal of our study was to provide a proof-of-concept validation of the anterior thalamic-hippocampal electrophysiological modulation for long-term improvement in memory function. A recent study reported lasting improvements in memory performance on a month timescale*(66)*. We found evidence for chronic enhancement of verbal memory on the scale of a year, which was reflected in biomarker neural activities. The theta activities induced during memory encoding were found to correlate session-by-session with longitudinal task performance. Previous longitudinal studies of anterior thalamic DBS effects on cognitive functions showed positive outcomes in neuropsychological assessments performed at relatively wide time intervals and with repetition-training effects*(53)*. Another longitudinal study of hippocampal DBS was also limited by the practice effects inherent to standard neuropsychological testing*(67)*. None of these employed electrophysiological biomarkers. Here, we were able to repeatedly test verbal memory on a weekly basis with concurrent LFP recordings in home environments. As a result, the anterior thalamic-hippocampal theta biomarkers of memory processing were demonstrated on a long time-scale of regularly repeated tests. This new chronic memory biomarker predicted the effect of therapeutic stimulation and general session performance. Hence, it can be used to quantify and track memory and cognitive processes across time, which may be more objective, robust and time-efficient than behavioral testing. It can also optimize the DBS therapy in an adaptive way over time to determine the parameters or targets of brain stimulation. Such simple biomarkers of induced spectral power are already applied in the adaptive DBS for movement disorders*(31)* and are feasible for implantable devices and brain computer interfaces to accelerate the development of therapeutic applications for modulating memory and higher brain functions.

## Materials and Methods

### Subjects

Four subjects with drug resistant epilepsy were implanted with the investigational Medtronic Summit RC+S™ Implantable Programmable Generator (IPG) with leads (model 3387 and 3391: 10.5 and 24 mm contact spacing, respectively) targeting bilateral anterior nucleus of the thalamus (ANT) and the hippocampus (Hc). The IPG provided continuous local field potential (LFP) time series data sampled at up to 500 Hz from four bipolar pairs selected from four out of sixteen leads. Time series data were remotely streamed to a cloud-based server accessible to clinician and researcher users*(25)*. This study was approved by the Food and Drug Administration (IDE G180224) and the Mayo Clinic Institutional Review Board. A fifth patient (M2) was consented but not implanted, and therefore is not included in this analysis. Patients received continuous or duty-cycle stimulation at either 2 Hz, 7 Hz, 100 Hz, or 145 Hz current frequency, within set ranges of pulse-width (90-200 us) and amplitude (1-6mA). Stimulation was halted immediately before scheduled performance of memory tasks to test modulatory effects of the chronic ANT stimulation, and then turned back on upon task completion to the same settings. No testing was performed around the time changing the stimulation parameters with at least one day of continuous chronic stimulation before any testing.

### Remote Delayed Free Recall Task

Patient cognitive abilities were probed using a previously described Free Recall Task *(39, 40, 54)*. The patients were able to complete verbal memory tasks in their home environments with LFP and behavioral data streamed to a handheld device and cloud repository. After the researcher initiated the task remotely, the participant was presented with up to 15 lists of words on a handheld tablet screen for a delayed test of free recall. Lists were composed of 12 randomly chosen nouns. Lists began with a 5 second countdown. Each word appeared on the screen for 1500 ms, with 1000 ms in between words without jitter. After a 20 s delay phase of simple addition equations answered with an onscreen keyboard, the participant was given 30 s to recall in any order as many of the words as possible recorded through a built-in microphone. Memory tasks were performed with synchronized ANT and Hc LFP data acquisition (<50 ms offset) that was artifact free due to halting stimulation for the duration of the testing. The researcher remained on the phone throughout the task.

### Power and Coherence Calculations

Hc and ANT power-in-band (PIB) measurements from the Medtronic Summit RC+S™ device were calculated by first downsampling the LFP signal by a factor of 5 for the theta band and a factor of 2 for the alpha band. For the beta and the broad-band gamma frequency bands, no downsampling was applied. Downsampling was applied to minimize inequalities in ratio between the sampling frequency and the upper cut-off frequency for the filter, as well as to shorten the computation time. Next, a linear trend was removed from the signal using Matlab “detrend” function (MathWorks Inc.), and the obtained signal was filtered for particular frequency bands, i.e., theta (4-8 Hz), alpha (9-13 Hz), beta (13-30 Hz), and broad-band gamma (30-120 Hz). An IIR elliptic filter with passband ripple of 0.1dB and stopband attenuation of 60dB was used to obtain the signals in the theta, alpha and beta frequency ranges. A Kaiser window FIR filter with the same passband ripple and stopband attenuation was used to obtain the signal in the broad-band gamma frequency range. The lower and upper stopband frequencies were equal to [3.37 Hz, 10.18 Hz], [7.56 Hz, 18.81 Hz], [10.96 Hz, 44.92 Hz], [25.29 Hz, 140.41 Hz], for the four frequency bands, respectively. The filter orders were equal to 14, 12, 14, 394, respectively for each frequency band. All the filters were automatically designed by the Matlab “bandpass” function (MathWorks Inc.), specific to the given frequency bands and the sampling frequency with consideration for the downsampling factor. For each filtered signal, a spectrogram was calculated using the Matlab “spectrogram” function (MathWorks Inc.) which applied the short-time Fourier transform with a window size *w*, specified by equation 1, where *fs* is a sampling frequency (equal to 500 Hz in the study) and *nd* is a downsampling factor. Window overlap was 95%.

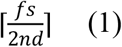

Next, the spectrogram was z-scored along its rows (each 1 Hz incremental bin). Columns of the spectrogram at indices *j* with visible filtering artifacts, resulting from non-physiological spikes and sharp-transitions, were removed from the analysis by applying a method specified by equations 2 and 3. *PSD* is a spectrogram of *n* rows by *m* columns, *Mdn* denotes a median function, and *thr* is a threshold value (equal to 18 in this study) for filtering out severe distortion and only non-physiological spikes.

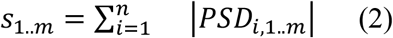

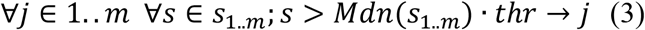

For each pair of filtered signals, acquired from different brain sites, a multi-taper time-frequency coherence in the form of a coherogram was calculated using the “cohgramc” function from the Chronux 2.11 toolbox (http://chronux.org/; *(68)*). For the theta, alpha, and beta signals 30 tapers were used and for the broad-band gamma signal 15 tapers were used. A time-bandwidth product parameter *TW* was equal to 1 in every case. The *movingwin* parameter was equal to [1 0.01]. All the analyses were performed in Matlab R2021b.

### Statistical Analysis

The Kruskal-Wallis test was used to determine statistical significance of the differences in mean spectral power or coherence in any of the groups (Fig. 2a: enc/delay/rec; Fig. 2b,c 3a,b: pre/on). In the case of the comparison between the encoding, delay, and recall periods, the null hypothesis was that the means from each patient data within each group came from the same distribution, i.e., the power/coherence was not modulated by the cognitive load associated with each of the three phases. The Tukey-Kramer post-hoc test was applied to determine statistically significant differences in mean power and coherence between pairs of enc./delay/rec. groups (Fig. 2a). In the case of the comparison between the pre/on periods, the null hypothesis was that the power/coherence means from each patient’s data was not modulated by the word or equation display event. For the word display event, the “pre” and “on” periods constituted 1s before and 1s after the word onset, respectively. For the equation display event, marked as 0s, the pre and on periods constituted a range of [-0.3s 0.8s] and (0.8s 3s], respectively. Those ranges were established based on the population-average threshold values across the spectrograms for all four sites. Spearman correlation was used to determine the correlation between the memory performance (number of recalled words) and power/coherence magnitudes in distinct brain sites and task periods, across consecutive sessions. Permutation test was applied to assess statistically significant differences in memory performance (mean number of recalled words) in consecutive sessions, between the groups of different stimulation frequencies (Fig. 4b, darker). 10,000 permutations were applied with randomized changing of labels for the stimulation session. The same approach was applied to assess statistical significance of the differences in mean power and coherence in all four brain sites and periods across consecutive sessions, between the groups of different stimulation frequencies (Fig. 4b, brighter). The significance level *α* in each case was equal to 0.05. All statistical tests were performed in Matlab R2021a. The permutation test was performed using an external GPL-3.0 function “permutationTest”*(69)*. Data and code available upon reasonable request.

## Supplementary Materials

**Supplementary Fig. S1**. Alpha band power changes during the task are dependent on brain area and phase of task.

**Supplementary Fig. S2**. Beta band power changes during the task are dependent on brain area and phase of task.

**Supplementary Fig. S3**. Gamma band power changes during the task are dependent on brain area and phase of task.

**Supplementary Fig. S4**. Population-average power and coherence changed during the equation display event.

**Supplementary Fig. S5**. Population-average alpha power and coherence changes during the word display event for recalled and forgotten words depended on brain area and recall state.

**Supplementary Fig. S6**. Population-average beta power and coherence during the word display event for recalled and forgotten words did not differ based on brain area or recall state.

**Supplementary Fig. S7**. Population-average gamma power and coherence during the word display event for recalled and forgotten words were dependent on brain area and recall state.

**Supplementary Fig. S8**. Performance of patients over time and theta power changed in respect to stimulation frequency.

## Acknowledgements

This research was supported by the National Institutes of Health (NIH; UH2&3-NS95495), by the First Team grant of the Foundation for Polish Science co-financed by the European Union under the European Regional Development Fund (Grant No. POIR.04.04.00-00-4379/17), and by the IDUB Aurum grant - Supporting International Research Team Building from the Gdansk University of Technology. V.S.M. designed the remote data collection protocol, collected data, and wrote the manuscript. M.L. analyzed data, made figures, and edited the manuscript. N.M.G., B.N.L., B.H.B., B.K., S.A.M., K.J.M., J.J.V, and G.A.W. are on the clinical teams for these patients and were responsible for decision making regarding electrode planning, surgical placement, medications, and stimulation settings. V.S., F.M., D.P.C., and V.K. were responsible for maintaining the streaming pipeline, tablet function, and data storage. J.D. designed the graphical user interface for the tablet-based tasks for the patients. E.A. provided insight on classical psychological interpretation of hippocampal-thalamic interactions. G.A.W. and M.T.K. were responsible for coordinating cross-institutional partnerships, providing funding, and editing the final manuscript. Medtronic provided the investigational Medtronic Summit RC+S™ devices.

## References and Notes

1. A. M. Lozano, N. Lipsman, H. Bergman, P. Brown, S. Chabardes, J. W. Chang, K. Matthews, C. C. McIntyre, T. E. Schlaepfer, M. Schulder, Y. Temel, J. Volkmann, J. K. Krauss, Deep brain stimulation: current challenges and future directions. Nat. Rev. Neurol. 15, 148–160 (2019).

2. B. J. Nuttin, L. A. Gabriëls, P. R. Cosyns, B. A. Meyerson, S. Andréewitch, S. G. Sunaert, A. F. Maes, P. J. Dupont, J. M. Gybels, F. Gielen, H. G. Demeulemeester, Long-term electrical capsular stimulation in patients with obsessive-compulsive disorder. Neurosurgery 62, 966–977 (2008).

3. D. Martinez-Ramirez, J. Jimenez-Shahed, J. F. Leckman, M. Porta, D. Servello, F.-G. Meng, J. Kuhn, D. Huys, J. C. Baldermann, T. Foltynie, M. I. Hariz, E. M. Joyce, L. Zrinzo, Z. Kefalopoulou, P. Silburn, T. Coyne, A. Y. Mogilner, M. H. Pourfar, S. M. Khandhar, M. Auyeung, J. L. Ostrem, V. Visser-Vandewalle, M.-L. Welter, L. Mallet, C. Karachi, J. L. Houeto, B. T. Klassen, L. Ackermans, T. Kaido, Y. Temel, R. E. Gross, H. C. Walker, A.M. Lozano, B. L. Walter, Z. Mari, W. S. Anderson, B. K. Changizi, E. Moro, S. E. Zauber, L. E. Schrock, J.-G. Zhang, W. Hu, K. Rizer, E. H. Monari, K. D. Foote, I. A. Malaty, W. Deeb, A. Gunduz, M. S. Okun, Efficacy and Safety of Deep Brain Stimulation in Tourette Syndrome: The International Tourette Syndrome Deep Brain Stimulation Public Database and Registry. JAMA Neurol. 75, 353–359 (2018).

4. P. E. Holtzheimer, C. Hamani, Deep brain stimulation for treatment-resistant depressionDeep Brain Stimulation: Technology and Applications (Volume 2), 64–75 (2014).

5. H. Mayberg, Deep brain stimulation for treatment-resistant depressionJournal of Affective Disorders 107, S23 (2008).

6. K. Kim, A. D. Ekstrom, N. Tandon, A network approach for modulating memory processes via direct and indirect brain stimulation: Toward a causal approach for the neural basis of memory. Neurobiol. Learn. Mem. 134 Pt A, 162–177 (2016).

7. N. Suthana, I. Fried, Deep brain stimulation for enhancement of learning and memory. Neuroimage 85 Pt 3, 996–1002 (2014).

8. N. Suthana, Z. M. Aghajan, E. A. Mankin, A. Lin, Reporting Guidelines and Issues to Consider for Using Intracranial Brain Stimulation in Studies of Human Declarative Memory. Front. Neurosci. 12, 905 (2018).

9. E. A. Mankin, I. Fried, Modulation of Human Memory by Deep Brain Stimulation of the Entorhinal-Hippocampal Circuitry. Neuron 106, 218–235 (2020).

10. E. L. Johnson, J. W. Y. Kam, A. Tzovara, R. T. Knight, Insights into human cognition from intracranial EEG: A review of audition, memory, internal cognition, and causality. J. Neural Eng. 17, 051001 (2020).

11. N. Suthana, Z. Haneef, J. Stern, R. Mukamel, E. Behnke, B. Knowlton, I. Fried, Memory enhancement and deep-brain stimulation of the entorhinal area. N. Engl. J. Med. 366, 502–510 (2012).

12. J. Jacobs, J. Miller, S. A. Lee, T. Coffey, A. J. Watrous, M. R. Sperling, A. Sharan, G. Worrell, B. Berry, B. Lega, B. C. Jobst, K. Davis, R. E. Gross, S. A. Sheth, Y. Ezzyat, S. R. Das, J. Stein, R. Gorniak, M. J. Kahana, D. S. Rizzuto, Direct Electrical Stimulation of the Human Entorhinal Region and Hippocampus Impairs Memory. Neuron 92, 983–990 (2016).

13. J.-M. S. Leoutsakos, H. Yan, W. S. Anderson, W. F. Asaad, G. Baltuch, A. Burke, M. M. Chakravarty, K. E. Drake, K. D. Foote, L. Fosdick, P. Giacobbe, Z. Mari, M. P. McAndrews, C. A. Munro, E. S. Oh, M. S. Okun, J. C. Pendergrass, F. A. Ponce, P. B. Rosenberg, M. N. Sabbagh, S. Salloway, D. F. Tang-Wai, S. D. Targum, D. Wolk, A. M. Lozano, G. S. Smith, C. G. Lyketsos, Deep Brain Stimulation Targeting the Fornix for Mild Alzheimer Dementia (the ADvance Trial): A Two Year Follow-up Including Results of Delayed Activation. J. Alzheimers. Dis. 64, 597–606 (2018).

14. A. M. Lozano, L. Fosdick, M. M. Chakravarty, J.-M. Leoutsakos, C. Munro, E. Oh, K. E. Drake, C. H. Lyman, P. B. Rosenberg, W. S. Anderson, D. F. Tang-Wai, J. C. Pendergrass, S. Salloway, W. F. Asaad, F. A. Ponce, A. Burke, M. Sabbagh, D. A. Wolk, G. Baltuch, M. S. Okun, K. D. Foote, M. P. McAndrews, P. Giacobbe, S. D. Targum, C. G. Lyketsos, G. S. Smith, A Phase II Study of Fornix Deep Brain Stimulation in Mild Alzheimer’s Disease. J. Alzheimers. Dis. 54, 777–787 (2016).

15. A. W. Laxton, D. F. Tang-Wai, M. P. McAndrews, D. Zumsteg, R. Wennberg, R. Keren, J. Wherrett, G. Naglie, C. Hamani, G. S. Smith, A. M. Lozano, A phase I trial of deep brain stimulation of memory circuits in Alzheimer’s disease. Ann. Neurol. 68, 521–534 (2010).

16. W. Deeb, B. Salvato, L. Almeida, K. D. Foote, R. Amaral, J. Germann, P. B. Rosenberg, D. F. Tang-Wai, D. A. Wolk, A. D. Burke, S. Salloway, M. N. Sabbagh, M. M. Chakravarty, G. S. Smith, C. G. Lyketsos, A. M. Lozano, M. S. Okun, Fornix-Region Deep Brain Stimulation-Induced Memory Flashbacks in Alzheimer’s Disease. N. Engl. J. Med. 381, 783–785 (2019).

17. D. Maltête, D. Wallon, J. Bourilhon, R. Lefaucheur, T. Danaila, S. Thobois, L. Defebvre, K. Dujardin, J.-L. Houeto, O. Godefroy, P. Krystkowiak, O. Martinaud, A. Gillibert, M. Chastan, P. Vera, D. Hannequin, M.-L. Welter, S. Derrey, Nucleus Basalis of Meynert Stimulation for Lewy Body DementiaNeurology 96, e684–e697 (2021).

18. W. Liu, D.-Y. Yu, Bilateral nucleus basalis of Meynert deep brain stimulation for dementia with Lewy bodies: A randomised clinical trialBrain Stimulation 13, 1612–1613 (2020).

19. W. Zhang, W. Liu, B. Patel, Y. Chen, K. Wang, A. Yang, F. Meng, A. Wagle Shukla, S. Cen, J. Yu, A. Ramirez-Zamora, J. Zhang, Case Report: Deep Brain Stimulation of the Nucleus Basalis of Meynert for Advanced Alzheimer’s Disease. Front. Hum. Neurosci. 15, 645584 (2021).

20. J. Kuhn, K. Hardenacke, E. Shubina, D. Lenartz, V. Visser-Vandewalle, K. Zilles, V. Sturm, H.-J. Freund, Deep Brain Stimulation of the Nucleus Basalis of Meynert in Early Stage of Alzheimer’s DementiaBrain Stimul. 8, 838–839 (2015).

21. J. Kuhn, K. Hardenacke, D. Lenartz, T. Gruendler, M. Ullsperger, C. Bartsch, J. K. Mai, K. Zilles, A. Bauer, A. Matusch, R.-J. Schulz, M. Noreik, C. P. Bührle, D. Maintz, C. Woopen, P. Häussermann, M. Hellmich, J. Klosterkötter, J. Wiltfang, M. Maarouf, H.-J. Freund, V. Sturm, Deep brain stimulation of the nucleus basalis of Meynert in Alzheimer’s dementia. Mol. Psychiatry 20, 353–360 (2015).

22. V. Sreekumar, J. H. Wittig Jr, T. C. Sheehan, K. A. Zaghloul, Principled Approaches to Direct Brain Stimulation for Cognitive Enhancement. Front. Neurosci. 11, 650 (2017).

23. S. Stanslaski, J. Herron, T. Chouinard, D. Bourget, B. Isaacson, V. Kremen, E. Opri, W. Drew, B. H. Brinkmann, A. Gunduz, T. Adamski, G. A. Worrell, T. Denison, A Chronically Implantable Neural Coprocessor for Investigating the Treatment of Neurological Disorders. IEEE Trans. Biomed. Circuits Syst. 12, 1230–1245 (2018).

24. V. Sladky, P. Nejedly, F. Mivalt, B. H. Brinkmann, I. Kim, E. K. St. Louis, N. M. Gregg, B. N. Lundstrom, C. M. Crowe, T. P. Attia, D. Crepeau, I. Balzekas, V. Marks, L. P. Wheeler, J. Cimbalnik, M. Cook, R. Janca, B. K. Sturges, K. Leyde, K. J. Miller, J. J. Van Gompel, T. Denison, G. A. Worrell, V. Kremen, Distributed brain co-processor for tracking electrophysiology and behavior during electrical brain stimulation, doi:10.1101/2021.03.08.434476.

25. V. Kremen, B. H. Brinkmann, I. Kim, H. Guragain, M. Nasseri, A. L. Magee, T. Pal Attia, P. Nejedly, V. Sladky, N. Nelson, S.-Y. Chang, J. A. Herron, T. Adamski, S. Baldassano, J. Cimbalnik, V. Vasoli, E. Fehrmann, T. Chouinard, E. E. Patterson, B. Litt, M. Stead, J. Van Gompel, B. K. Sturges, H. J. Jo, C. M. Crowe, T. Denison, G. A. Worrell, Integrating Brain Implants With Local and Distributed Computing Devices: A Next Generation Epilepsy Management System. IEEE J Transl Eng Health Med 6, 2500112 (2018).

26. T. P. Attia, D. Crepeau, V. Kremen, M. Nasseri, H. Guragain, S. W. Steele, V. Sladky, P. Nejedly, F. Mivalt, J. A. Herron, M. Stead, T. Denison, G. A. Worrell, B. H. Brinkmann, Epilepsy Personal Assistant Device—A Mobile Platform for Brain State, Dense Behavioral and Physiology Tracking and Controlling Adaptive StimulationFrontiers in Neurology 12 (2021), doi:10.3389/fneur.2021.704170.

27. F. Mivalt, V. Kremen, V. Sladky, I. Balzekas, P. Nejedly, N. Gregg, B. Lundstrom, K. Lepkova, T. Pridalova, B. H. Brinkmann, P. Jurak, J. J. Van Gompel, K. Miller, T. Denison, E. St Louis, G. A. Worrell, Electrical Brain Stimulation and Continuous Behavioral State Tracking in Ambulatory Humans, doi:10.1101/2021.08.10.21261645.

28. G. A. Worrell, Electrical Brain Stimulation for Epilepsy and Emerging ApplicationsJournal of Clinical Neurophysiology 38, 471–477 (2021).

29. N. M. Gregg, V. Sladky, P. Nejedly, F. Mivalt, I. Kim, I. Balzekas, B. K. Sturges, C. Crowe, E. E. Patterson, J. J. Van Gompel, B. N. Lundstrom, K. Leyde, T. J. Denison, B. H. Brinkmann, V. Kremen, G. A. Worrell, Thalamic deep brain stimulation modulates cycles of seizure risk in epilepsy. Sci. Rep. 11, 24250 (2021).

30. N. M. Gregg, V. S. Marks, V. Sladky, B. N. Lundstrom, B. Klassen, S. A. Messina, B. H. Brinkmann, K. J. Miller, J. J. Van Gompel, V. Kremen, G. A. Worrell, Anterior nucleus of the thalamus seizure detection in ambulatory humans. Epilepsia 62, e158–e164 (2021).

31. R. ‘ee Gilron, S. Little, R. Perrone, R. Wilt, C. de Hemptinne, M. S. Yaroshinsky, C. A. Racine, S. S. Wang, J. L. Ostrem, P. S. Larson, D. D. Wang, N. B. Galifianakis, I. O. Bledsoe, M. San Luciano, H. E. Dawes, G. A. Worrell, V. Kremen, D. A. Borton, T. Denison, P. A. Starr, Long-term wireless streaming of neural recordings for circuit discovery and adaptive stimulation in individuals with Parkinson’s disease. Nat. Biotechnol. 39, 1078–1085 (2021).

32. N. C. Swann, C. de Hemptinne, S. Miocinovic, S. Qasim, J. L. Ostrem, N. B. Galifianakis, M. S. Luciano, S. S. Wang, N. Ziman, R. Taylor, P. A. Starr, Chronic multisite brain recordings from a totally implantable bidirectional neural interface: experience in 5 patients with Parkinson’s disease. J. Neurosurg. 128, 605–616 (2018).

33. S. Little, E. Tripoliti, M. Beudel, A. Pogosyan, H. Cagnan, D. Herz, S. Bestmann, T. Aziz, B. Cheeran, L. Zrinzo, M. Hariz, J. Hyam, P. Limousin, T. Foltynie, P. Brown, Adaptive deep brain stimulation for Parkinson’s disease demonstrates reduced speech side effects compared to conventional stimulation in the acute setting. J. Neurol. Neurosurg. Psychiatry 87, 1388–1389 (2016).

34. A. C. Meidahl, G. Tinkhauser, D. M. Herz, H. Cagnan, J. Debarros, P. Brown, Adaptive Deep Brain Stimulation for Movement Disorders: The Long Road to Clinical TherapyMovement Disorders 32, 810–819 (2017).

35. S. Little, A. Pogosyan, S. Neal, B. Zavala, L. Zrinzo, M. Hariz, T. Foltynie, P. Limousin, K. Ashkan, J. FitzGerald, A. L. Green, T. Z. Aziz, P. Brown, Adaptive deep brain stimulation in advanced Parkinson disease. Ann. Neurol. 74, 449–457 (2013).

36. V. Kremen, B. H. Brinkmann, J. J. Van Gompel, M. Stead, E. K. St Louis, G. A. Worrell, Automated unsupervised behavioral state classification using intracranial electrophysiologyJournal of Neural Engineering 16, 026004 (2019).

37. Y. Ezzyat, J. E. Kragel, J. F. Burke, D. F. Levy, A. Lyalenko, P. Wanda, L. O’Sullivan, K. B. Hurley, S. Busygin, I. Pedisich, M. R. Sperling, G. A. Worrell, M. T. Kucewicz, K. A. Davis, T. H. Lucas, C. S. Inman, B. C. Lega, B. C. Jobst, S. A. Sheth, K. Zaghloul, M. J. Jutras, J. M. Stein, S. R. Das, R. Gorniak, D. S. Rizzuto, M. J. Kahana, Direct Brain Stimulation Modulates Encoding States and Memory Performance in HumansCurrent Biology 27, 1251–1258 (2017).

38. K. V. Saboo, Y. Varatharajah, B. M. Berry, M. R. Sperling, R. Gorniak, K. A. Davis, B. C. Jobst, R. E. Gross, B. Lega, S. A. Sheth, M. J. Kahana, M. T. Kucewicz, G. A. Worrell, R. K. Iyer, A Computationally Efficient Model for Predicting Successful Memory Encoding Using Machine-Learning-based EEG Channel Selection2019 9th International IEEE/EMBS Conference on Neural Engineering (NER) (2019), doi:10.1109/ner.2019.8717057.

39. V. S. Marks, K. V. Saboo, Ç. Topçu, M. Lech, T. P. Thayib, P. Nejedly, V. Kremen, G. A. Worrell, M. T. Kucewicz, Independent dynamics of low, intermediate, and high frequency spectral intracranial EEG activities during human memory formation. Neuroimage 245, 118637 (2021).

40. Ç. Topçu, V. S. Marks, K. V. Saboo, M. Lech, P. Nejedly, V. Kremen, G. A. Worrell, M. T. Kucewicz, Hotspot of human verbal memory encoding in the left anterior prefrontal cortexeBioMedicine 82, 104135 (2022).

41. Y. Ezzyat, P. A. Wanda, D. F. Levy, A. Kadel, A. Aka, I. Pedisich, M. R. Sperling, A. D. Sharan, B. C. Lega, A. Burks, R. E. Gross, C. S. Inman, B. C. Jobst, M. A. Gorenstein, K. A. Davis, G. A. Worrell, M. T. Kucewicz, J. M. Stein, R. Gorniak, S. R. Das, D. S. Rizzuto, M. J. Kahana, Closed-loop stimulation of temporal cortex rescues functional networks and improves memory. Nat. Commun. 9, 365 (2018).

42. M. T. Kucewicz, B. M. Berry, L. R. Miller, F. Khadjevand, Y. Ezzyat, J. M. Stein, V. Kremen, B. H. Brinkmann, P. Wanda, M. R. Sperling, R. Gorniak, K. A. Davis, B. C. Jobst, R. E. Gross, B. Lega, J. Van Gompel, S. M. Stead, D. S. Rizzuto, M. J. Kahana, G. A. Worrell, Evidence for verbal memory enhancement with electrical brain stimulation in the lateral temporal cortex. Brain 141, 971–978 (2018).

43. N. D. Child, E. E. Benarroch, Anterior nucleus of the thalamus: Functional organization and clinical implicationsNeurology 81, 1869–1876 (2013).

44. R. Fisher, V. Salanova, T. Witt, R. Worth, T. Henry, R. Gross, K. Oommen, I. Osorio, J. Nazzaro, D. Labar, M. Kaplitt, M. Sperling, E. Sandok, J. Neal, A. Handforth, J. Stern DeSalles, S. Chung, A. Shetter, D. Bergen, R. Bakay, J. Henderson, J. French, G. Baltuch, W. Rosenfeld, A. Youkilis, W. Marks, P. Garcia, N. Barbaro, N. Fountain, C. Bazil, R. Goodman, G. McKhann, K. Babu Krishnamurthy, S. Papavassiliou, C. Epstein, J. Pollard, L. Tonder, J. Grebin, R. Coffey, N. Graves, the SANTE Study Group, Electrical stimulation of the anterior nucleus of thalamus for treatment of refractory epilepsyEpilepsia 51, 899–908 (2010).

45. A. J. D. Nelson, The anterior thalamic nuclei and cognition: A role beyond space? Neurosci. Biobehav. Rev. 126, 1–11 (2021).

46. M. Leszczyński, T. Staudigl, Memory-guided attention in the anterior thalamusNeuroscience & Biobehavioral Reviews 66, 163–165 (2016).

47. J. P. Aggleton, S. M. O’Mara, The anterior thalamic nuclei: core components of a tripartite episodic memory system. Nat. Rev. Neurosci. 23, 505–516 (2022).

48. C. M. Sweeney-Reed, L. Buentjen, J. Voges, F. C. Schmitt, T. Zaehle, J. W. Y. Kam, J. Kaufmann, H.-J. Heinze, H. Hinrichs, R. T. Knight, M. D. Rugg, The role of the anterior nuclei of the thalamus in human memory processing. Neurosci. Biobehav. Rev. 126, 146– 158 (2021).

49. K. Štillová, P. Jurák, J. Chládek, J. Chrastina, J. Halámek, M. Bočková, S. Goldemundová, I. Říha, I. Rektor, The Role of Anterior Nuclei of the Thalamus: A Subcortical Gate in Memory Processing: An Intracerebral Recording Study. PLoS One 10, e0140778 (2015).

50. D. S. Roy, Y. Zhang, T. Aida, C. Shen, K. M. Skaggs, Y. Hou, M. Fleishman, O. Mosto, A. Weninger, G. Feng, Anterior thalamic circuits crucial for working memory. Proc. Natl. Acad. Sci. U. S. A. 119, e2118712119 (2022).

51. S. Jun, J. S. Kim, C. K. Chung, Direct Stimulation of Human Hippocampus During Verbal Associative Encoding Enhances Subsequent Memory Recollection. Front. Hum. Neurosci. 13, 23 (2019).

52. Y.-S. Oh, H. J. Kim, K. J. Lee, Y. I. Kim, S.-C. Lim, Y.-M. Shon, Cognitive improvement after long-term electrical stimulation of bilateral anterior thalamic nucleus in refractory epilepsy patients. Seizure 21, 183–187 (2012).

53. A. I. Tröster, K. J. Meador, C. P. Irwin, R. S. Fisher, SANTE Study Group, Memory and mood outcomes after anterior thalamic stimulation for refractory partial epilepsy. Seizure 45, 133–141 (2017).

54. V. S. Marks, T. J. Richner, N. M. Gregg, V. Sladky, J. Dolezal, V. Kremen, G. A. Worrell, M. T. Kucewicz, Deep Brain Stimulation of Anterior Nuclei of the Thalamus and Hippocampal Seizure Rate Modulate Verbal Memory Performance2022 IEEE International Conference on Electro Information Technology (eIT) (2022), doi:10.1109/eit53891.2022.9813930.

55. M. D. Rugg, Memories Are Made of This. Science 281, 1151–1152 (1998).

56. M. Stangl, U. Topalovic, C. S. Inman, S. Hiller, D. Villaroman, Z. M. Aghajan, L. Christov-Moore, N. R. Hasulak, V. R. Rao, C. H. Halpern, D. Eliashiv, I. Fried, N. Suthana, Boundary-anchored neural mechanisms of location-encoding for self and others. Nature 589, 420–425 (2021).

57. I. Balzekas, V. Sladky, N. Gregg, F. Mivalt, V. Marks, B. Lundstrom, J. Van Gompel, V. Kremen, B. Brinkmann, G. Worrell, Hippocampal-ANT connectivity and ANT DBS: Circadian trends and response to stimulationBrain Stimulation 14, 1654 (2021).

58. I. Balzekas, V. Sladky, P. Nejedly, B. H. Brinkmann, D. Crepeau, F. Mivalt, N. M. Gregg, T. Pal Attia, V. S. Marks, L. Wheeler, T. E. Riccelli, J. P. Staab, B. N. Lundstrom, K. J. Miller, J. Van Gompel, V. Kremen, P. E. Croarkin, G. A. Worrell, Invasive Electrophysiology for Circuit Discovery and Study of Comorbid Psychiatric Disorders in Patients With Epilepsy: Challenges, Opportunities, and Novel Technologies. Front. Hum. Neurosci. 15 (2021), doi:10.3389/fnhum.2021.702605.

59. L. Sun, J. Peräkylä, M. Polvivaara, J. Öhman, J. Peltola, K. Lehtimäki, H. Huhtala, K. M. Hartikainen, Human anterior thalamic nuclei are involved in emotion–attention interaction. Neuropsychologia 78, 88–94 (2015).

60. A. Goyal, J. Miller, S. E. Qasim, A. J. Watrous, J. M. Stein, C. S. Inman, R. E. Gross, J. T. Willie, B. Lega, J.-J. Lin, A. Sharan, C. Wu, M. R. Sperling, S. A. Sheth, G. M. McKhann, E. H. Smith, C. Schevon, J. Jacobs, Functionally distinct high and low theta oscillations in the human hippocampus, doi:10.1101/498055.

61. Z. J. Urgolites, J. T. Wixted, S. D. Goldinger, M. H. Papesh, D. M. Treiman, L. R. Squire, P. N. Steinmetz, Spiking activity in the human hippocampus prior to encoding predicts subsequent memory. Proc. Natl. Acad. Sci. U. S. A. 117, 13767–13770 (2020).

62. C. M. Sweeney-Reed, T. Zaehle, J. Voges, F. C. Schmitt, L. Buentjen, K. Kopitzki, C. Esslinger, H. Hinrichs, H.-J. Heinze, R. T. Knight, A. Richardson-Klavehn, Corticothalamic phase synchrony and cross-frequency coupling predict human memory formationeLife 3 (2014), doi:10.7554/elife.05352.

63. C. M. Sweeney-Reed, T. Zaehle, J. Voges, F. C. Schmitt, L. Buentjen, K. Kopitzki, A. Richardson-Klavehn, H. Hinrichs, H.-J. Heinze, R. T. Knight, M. D. Rugg, Pre-stimulus thalamic theta power predicts human memory formation. Neuroimage 138, 100–108 (2016).

64. J. Liu, T. Yu, J. Wu, Y. Pan, Z. Tan, R. Liu, X. Wang, L. Ren, L. Wang, Anterior thalamic stimulation improves working memory precision judgments. Brain Stimul. 14, 1073– 1080 (2021).

65. R. J. Piper, R. M. Richardson, G. Worrell, D. W. Carmichael, T. Baldeweg, B. Litt, T. Denison, M. M. Tisdall, Towards network-guided neuromodulation for epilepsy. Brain (2022), doi:10.1093/brain/awac234.

66. S. Grover, W. Wen, V. Viswanathan, C. T. Gill, R. M. G. Reinhart, Long-lasting, dissociable improvements in working memory and long-term memory in older adults with repetitive neuromodulation. Nat. Neurosci. (2022), doi:10.1038/s41593-022-01132-3.

67. G. K. Bergey, M. J. Morrell, E. M. Mizrahi, A. Goldman, D. King-Stephens, D. Nair, S. Srinivasan, B. Jobst, R. E. Gross, D. C. Shields, G. Barkley, V. Salanova, P. Olejniczak, a, Cole, S. S. Cash, K. Noe, R. Wharen, G. Worrell, A. M. Murro, J. Edwards, M. Duchowny, D. Spencer, M. Smith, E. Geller, R. Gwinn, C. Skidmore, S. Eisenschenk, M. Berg, C. Heck, P. Van Ness, N. Fountain, P. Rutecki, A. Massey, C. O’Donovan, D. Labar, R. B. Duckrow, L. J. Hirsch, T. Courtney, F. T. Sun, C. G. Seale, Long-term treatment with responsive brain stimulation in adults with refractory partial seizures. Neurology 84, 810–817 (2015).

68. P. Mitra, Observed Brain Dynamics (Oxford University Press, 2007).

69. L. Krol, Permutation test (2022; https://github.com/lrkrol/permutationTest).

