## Supplemental figures for "Chronic modulation of human memory and thalamic-hippocampal theta activities"

Marks V.S. et al.

Supplementary Information

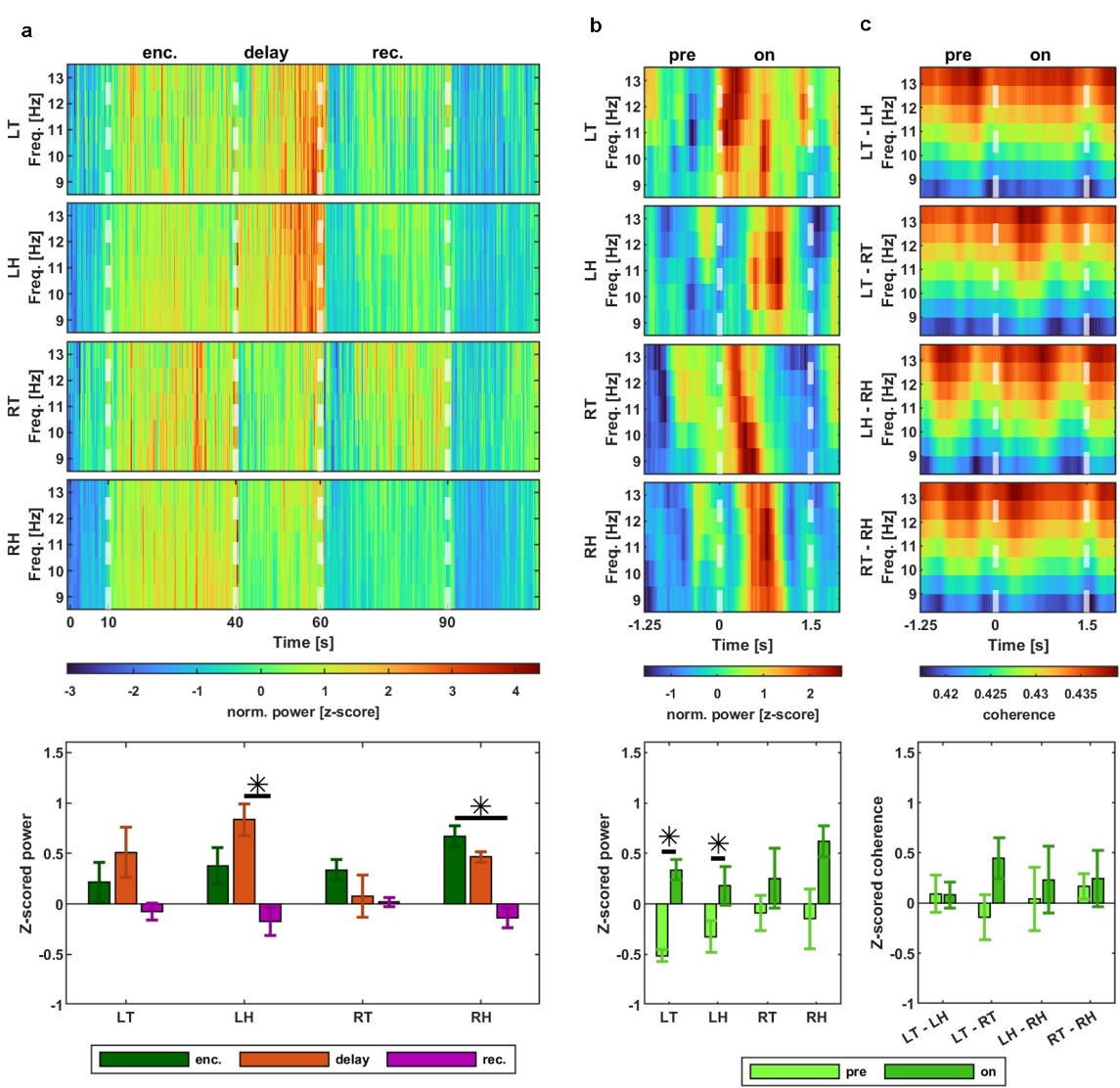

Supplementary Fig. S1. Alpha band power changes during the task are dependent on brain area and phase of task. Population-average alpha power across encoding, delay and recall phases (a) and then power (b) and coherence (c) specifically during the word display event.

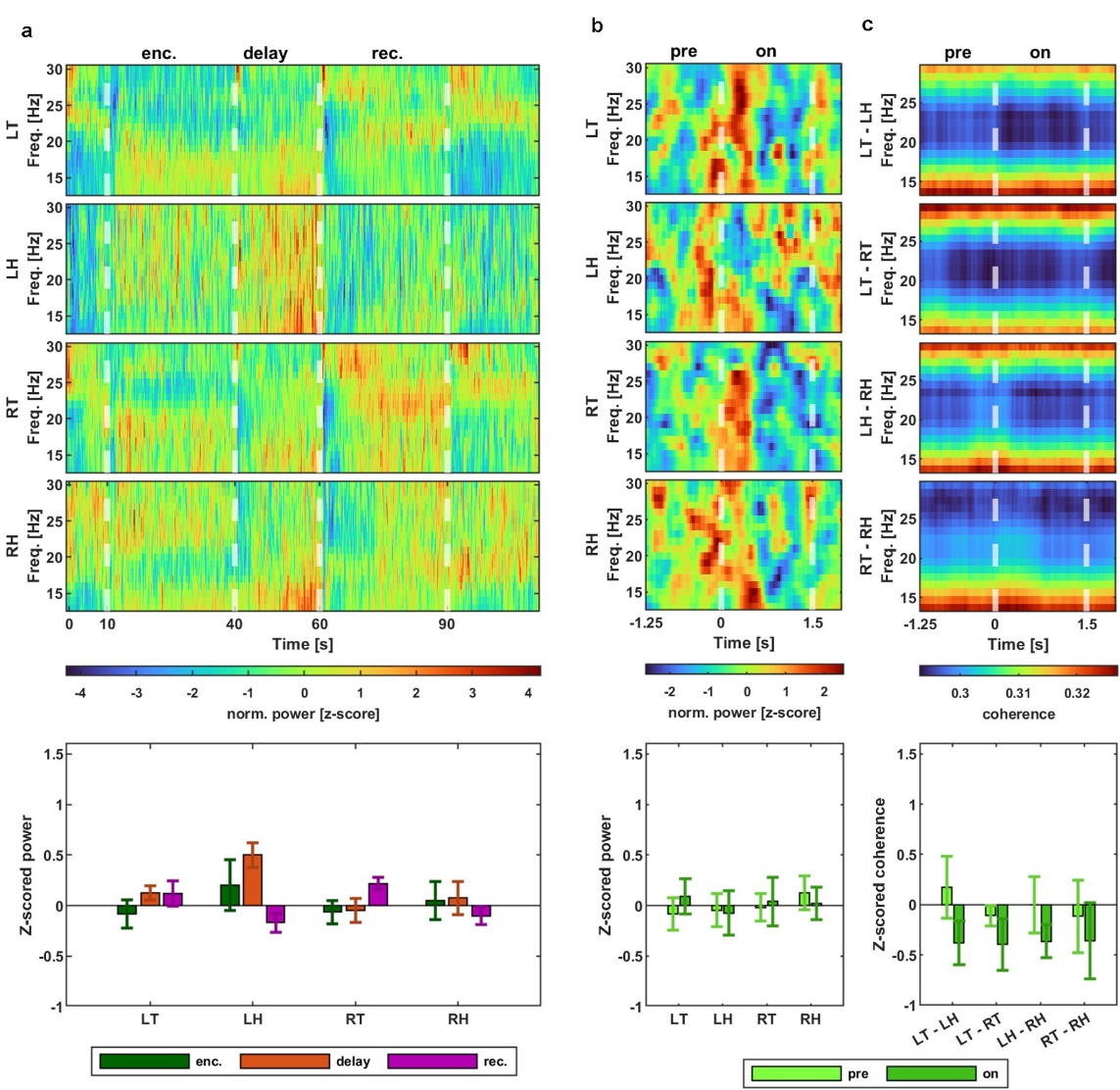

Supplementary Fig. S2. Beta band power changes during the task are dependent on brain area and phase of task. Population-average beta power across encoding, delay and recall phases (a) and then power (b) and coherence (c) specifically during the word display event.

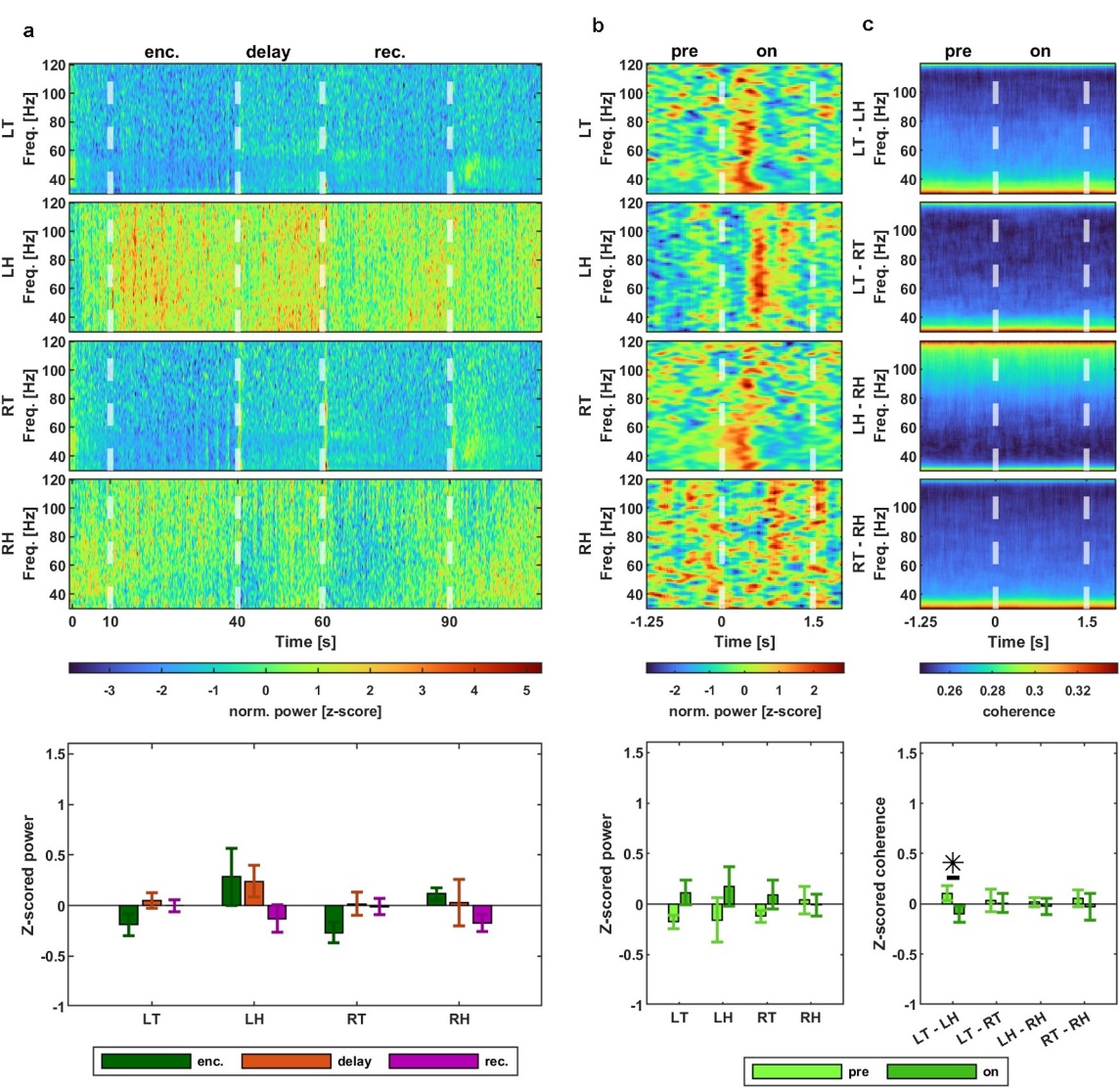

Supplementary Fig. S3. Gamma band power changes during the task are dependent on brain area and phase of task. Population-average gamma power across encoding, delay and recall phases (a) and then power (b) and coherence (c) specifically during the word display event.

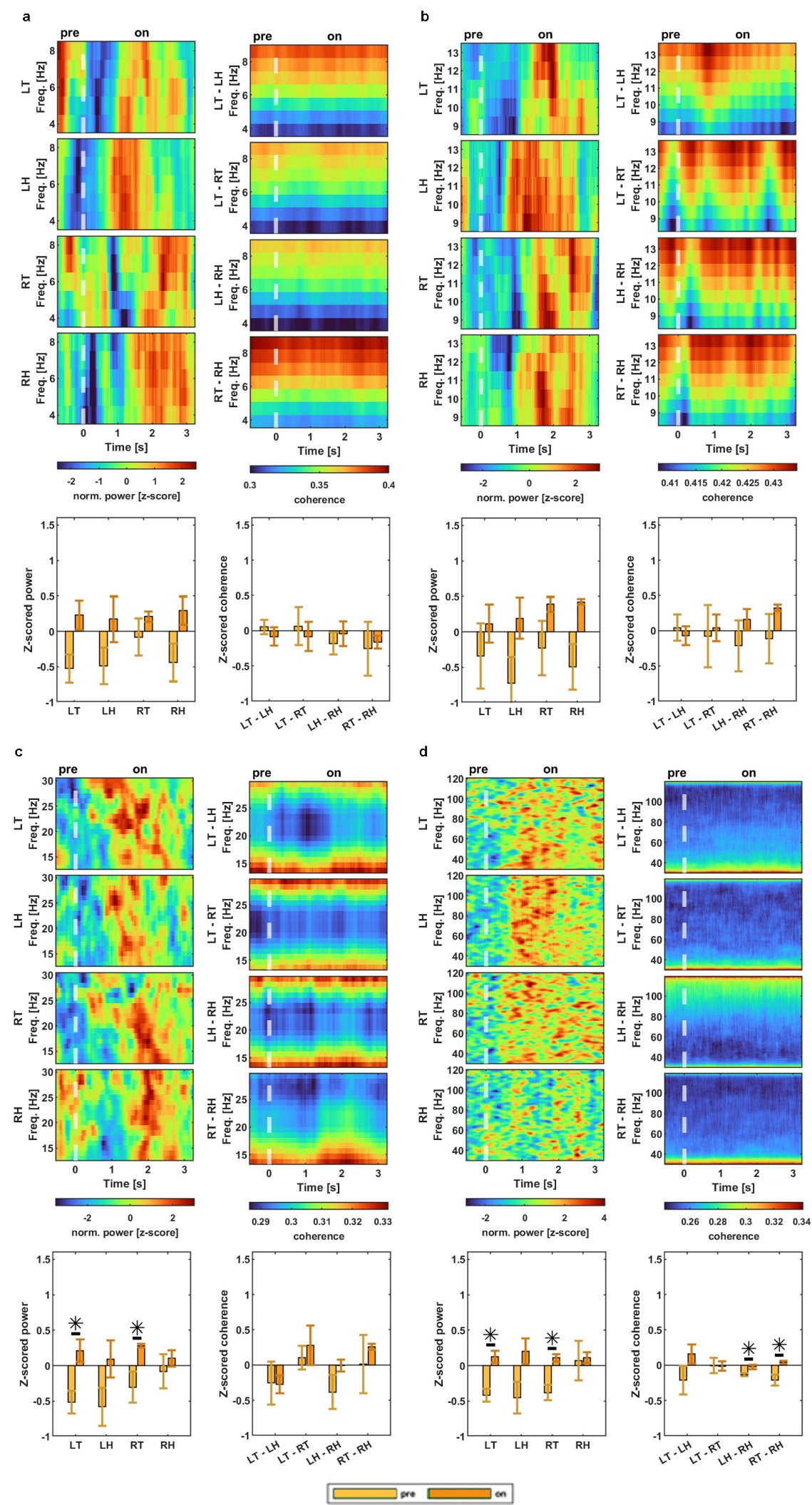

Supplementary Fig. S4. Population-average power and coherence changed during the equation display event. Theta (a), alpha (b), beta (c), and gamma (d) frequency bands displayed.

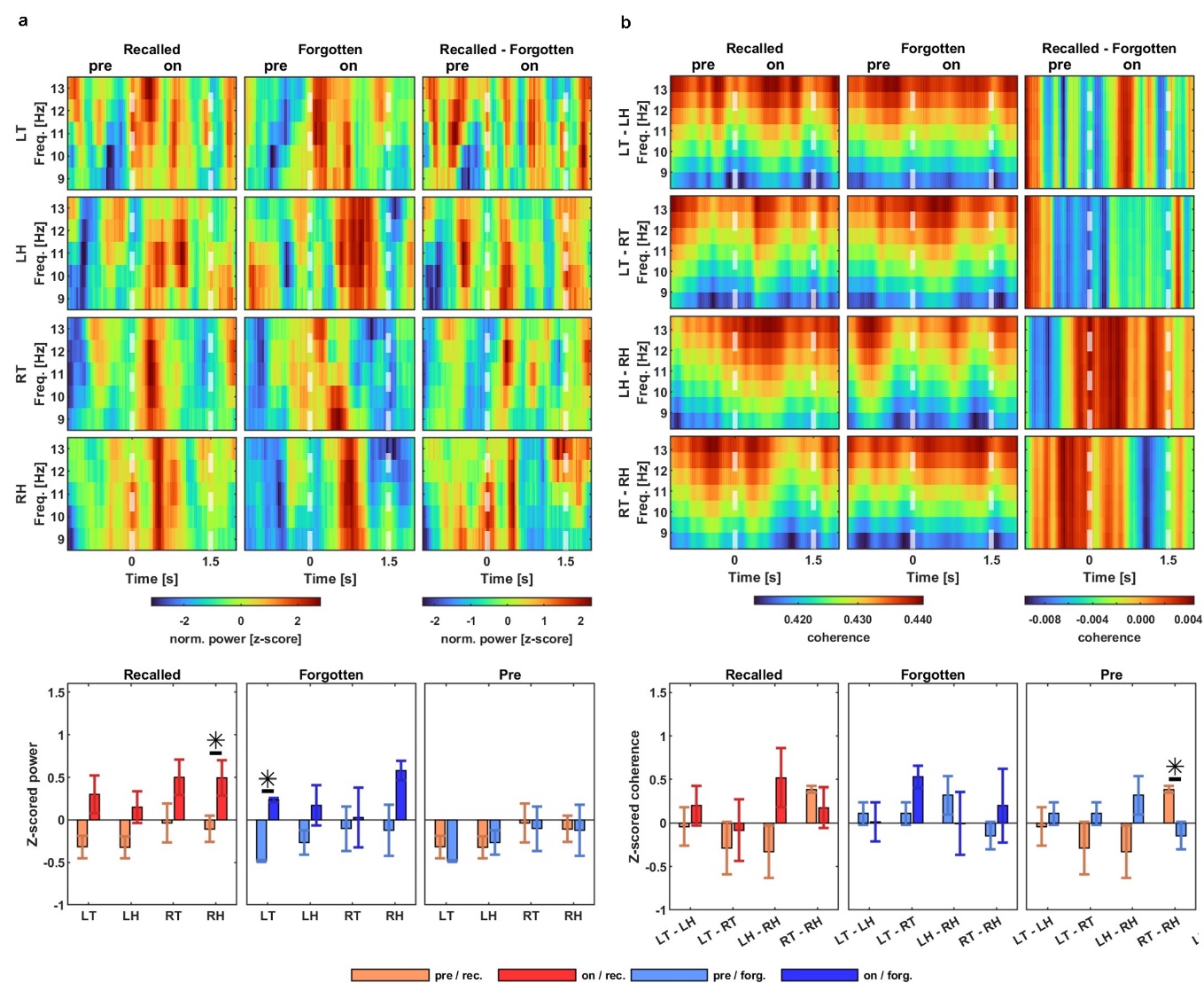

Supplementary Fig. S5. Population-average alpha power and coherence changes during the word display event for recalled and forgotten words depended on brain area and recall state.

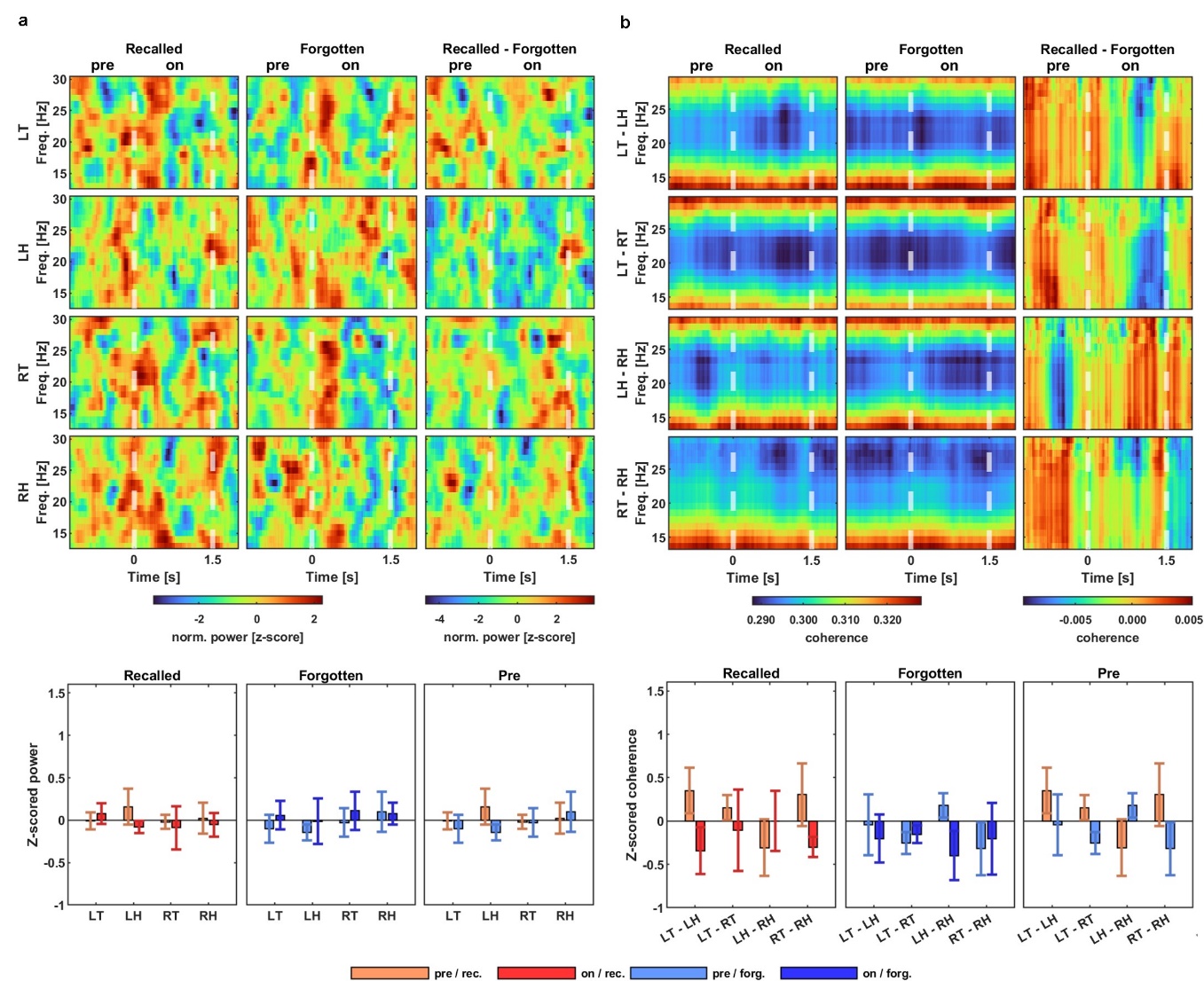

Supplementary Fig. S6. Population-average beta power and coherence during the word display event for recalled and forgotten words did not differ based on brain area or recall state.

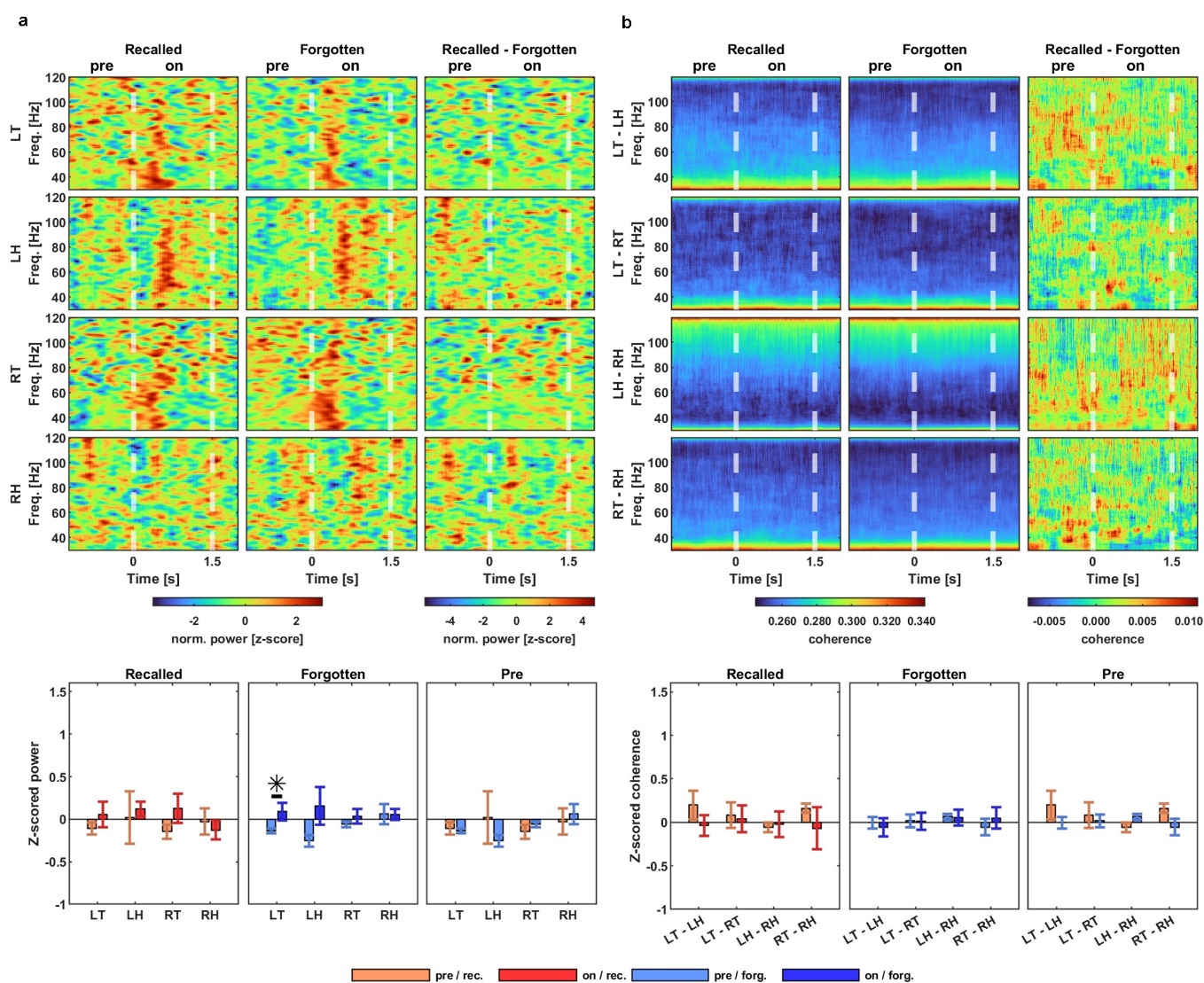

Supplementary Fig. S7. Population-average gamma power and coherence during the word display event for recalled and forgotten words were dependent on brain area and recall state.

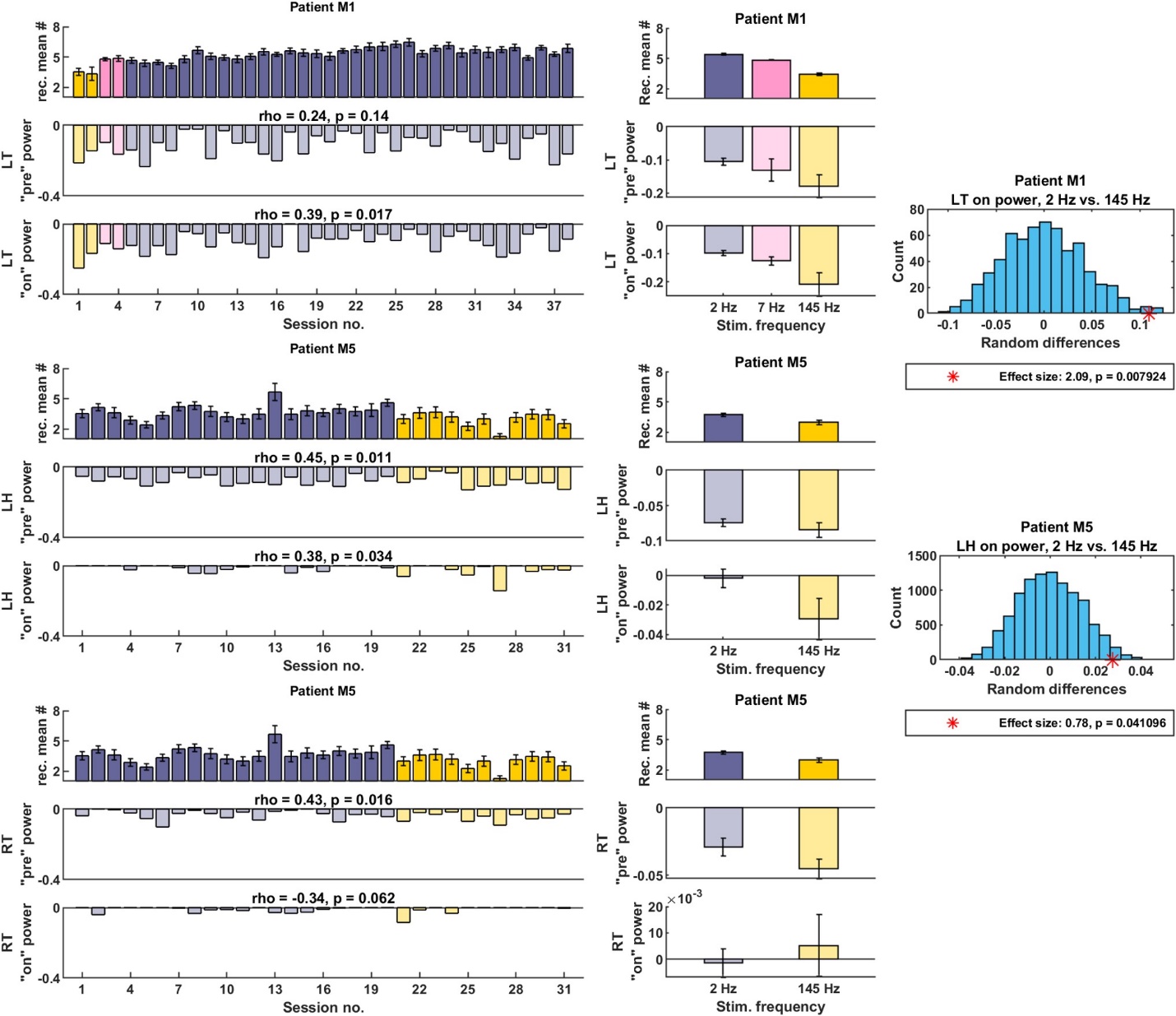

Supplementary Fig. S8. Performance of patients over time and theta power changed in respect to stimulation frequency.
